# Genomes of *Leishmania* parasites directly sequenced from patients with visceral leishmaniasis in the Indian subcontinent

**DOI:** 10.1101/676163

**Authors:** Malgorzata A. Domagalska, Hideo Imamura, Mandy Sanders, Frederik Van den Broeck, Narayan Raj Bhattarai, Manu Vanaerschot, Ilse Maes, Erika D’Haenens, Keshav Rai, Suman Rijal, Matthew Berriman, James A. Cotton, Jean-Claude Dujardin

## Abstract

Whole genome sequencing (WGS) is increasingly used for molecular diagnosis and epidemiology of infectious diseases. Current *Leishmania* genomic studies rely on DNA extracted from cultured parasites, which might introduce sampling and biological biases into the subsequent analyses. Up to now, direct analysis of *Leishmania* genome in clinical samples is hampered by high levels of human DNA and large variation in parasite load in patient samples. Here, we present a method, based on target enrichment of *Leishmania donovani* DNA with Agilent SureSelect technology, that allows the analysis of *Leishmania* genomes directly in clinical samples. We validated our protocol with a set of artificially mixed samples, followed by the analysis of 63 clinical samples (bone marrow or spleen aspirates) from visceral leishmaniasis patients in Nepal. We were able to identify genotypes using a set of diagnostic SNPs in almost all of these samples (97%) and access comprehensive genome-wide information in most (83%). This allowed us to perform phylogenomic analysis, assess chromosome copy number and identify large copy number variants (CNVs). Pairwise comparisons between the parasite genomes in clinical samples and derived *in vitro* cultured promastigotes showed a lower aneuploidy in amastigotes as well as genomic differences, suggesting polyclonal infections in patients. Altogether our results underline the need for sequencing parasite genomes directly in the host samples.

**Author summary:** Visceral leishmaniasis (VL) is caused by parasitic protozoa of the *Leishmania donovani* complex and is lethal in the absence of treatment. Whole Genome Sequencing (WGS) of *L. donovani* clinical isolates revealed hitherto cryptic population structure in the Indian Sub-Continent and provided insights into the epidemiology and potential mechanisms of drug resistance. However, several biases are likely introduced during the culture step. We report here the development of a method that allows determination of parasite genomes directly in clinical samples, and validate it on bone marrow and splenic aspirates of VL patients in Nepal. Our study sheds a new light on the biology of Leishmania in the human host: we found that intracellular parasites of the patients had very low levels of aneuploidy, in sharp contrast to the situation in cultivated isolates. Moreover, the observed differences in genomes between intracellular amastigotes of the patient and the derived cultured parasites suggests polyclonality of infections, with different clones dominating in clinical samples and in culture, likely due to fitness differences. We believe this method is most suitable for clinical studies and for molecular tracking in the context of elimination programs.

## Introduction

*Leishmania* (Kinetoplastida, Trypanosomatidae) is a genus of parasitic protozoa transmitted by sand flies to a variety of mammal hosts including humans. Within mammals, the amastigote-stage parasites are strictly intracellular and infect a range of professional phagocytes. *Leishmania* cause a broad spectrum of clinical presentations, the most severe being visceral leishmaniasis (VL) which annually affects up to 90,000 new individuals [1] and has a 10% mortality rate [2]. Ongoing control programs, like the one running in the Indian subcontinent (ISC) [3] could be challenged by several factors, including drug resistance [4, 5] and emergence of new parasite variants [6, 7]. More than ever, close surveillance is required to avoid new epidemics, including molecular tracking of the etiological agent.

Whole genome sequencing (WGS) is increasingly used for highly discriminatory parasite molecular tracking. Using that method, we previously resolved the hitherto cryptic population structure of *Leishmania donovani*, the etiological agent of visceral leishmaniasis (VL), in the Indian subcontinent ISC [8]. We identified a core group (CG) – associated with the successive VL epidemics in the lowlands – itself structured into (i) six congruent monophyletic groups (ISC2-7) together with (ii) three other groups (ISC8-10) and several ungrouped isolates, with a less certain evolutionary history. We found in the Nepalese highlands an emerging population (ISC1), quite different from the CG [6, 9]. We described an unprecedented level of aneuploidy and found genetic variants likely to underpin reduced efficacy of antimonial drugs in that region.

However, such studies suffer a major limitation: they currently rely on sequencing parasites isolated from patients and grown axenically *in vitro*, with two possible biases. First, there could be a major representation bias, because of the variable isolation success and the high frequency (up to 90%) of asymptomatic cases [10], currently unsampled. Second, we have shown experimentally major differences in the genome of *L. donovani* during the life cycle: in particular, aneuploidy was much lower and affected other chromosomes in the intra-cellular life stages (amastigotes) of infected hamsters, in comparison to cultivated promastigotes [11]. This genome plasticity may affect gene dosage and therefore have a biological impact via transcriptome and proteome changes [12]. If the phenomenon also occurs between amastigotes from human subjects and *in vitro* cultivated promastigotes, it could have a major impact on the search for molecular bio-markers for clinically important traits such as drug resistance or virulence, as relevant variants present in the host could be diluted or disappear in culture.

Applying parasite WGS directly in host samples thus appears critical for next generation genetic analyses of natural populations of *Leishmania*. Such applications are challenged by the low amounts of *Leishmania* DNA in biological samples and the high abundance of DNA in the nucleated host cells. Accordingly, methods for enrichment of parasite DNA are required before undertaking WGS. A series of approaches have been used for other pathogens, including (i) removal of methylated host DNA [13], (ii) selective whole genome amplification [14] or (iii) targeted genome enrichment [15]. Here, we explored the last solution. Using Agilent SureSelect technology, we designed an array for application in the *L. donovani* species complex. In this method an Illumina-based NGS sequencing library is prepared from the mixed genomic DNA sample, which is later hybridized with biotinylated RNA baits complementary to the *Leishmania* genome and the fraction of the input DNA that anneals to the baits is then sequenced (further called SuSL-seq). We validated and optimized this genome capture procedure with artificial mixtures of *Leishmania* and host DNA to simulate the proportions expected in clinical samples. Finally, we applied the method on a set of 63 bone marrow and splenic aspirates from Nepalese VL patients. In 12 of them, we compared the karyotype and genome sequence with that of isolates derived from the same tissue samples. We find genomic differences between the parasites analyzed directly in clinical samples and the derived isolates, further supporting the need for direct analyses in host samples.

## Methods

### Ethics statement

The ethics committee of the Nepal Health Research Council, Kathmandu (IRC/637/014) and the corresponding bodies at the Institute of Tropical Medicine of Antwerp and the Antwerp University, Belgium (B300201627694), reviewed and approved the study protocol and the use of already-existing samples. Informed written consent was obtained from each patient or his/her guardian for those <18 years of age. All the patients and caretakers/parents had the study purpose explained to them in local language. A total of 63 anonymized samples (59 bone marrows, 4 spleen aspirates from clinically confirmed VL patients in Terai, Nepal) were collected at the B.P. Koirala Institute of Health Sciences (BPKIHS) in Dharan, Nepal in the frame of four previous projects: (i) VL-project, sampling between 2000 and 2001, (ii) Leishnatdrug-R, sampling between 2002 and 2003, (iii) Kaladrug-R, sampling between 2009 and 2010 and (iv) SINGLE, sampling between 2014 and 2015. These samples are part of the same clinical studies that generated parasite isolates analyzed in [8].

### DNA extraction and next generation sequencing

DNA from parasite cultures and clinical samples (bone marrow and spleen aspirates) was extracted using QiaAmp DNA blood mini kit (Qiagen, Venlo, the Netherlands) following the manufacturer’s instructions, and DNA concentrations were verified using the Qubit broad-range DNA quantification kit (Thermo Fisher Scientific, Waltham, USA). Sequencing was done following the manufacturer’s standard cluster generation and sequencing protocols

<u>Standard Whole Genome Sequencing</u> was applied to isolates. Genomic DNA was sheared into 400bp fragments by focused ultrasonication (Covaris Inc., Woburn, USA). Amplification-free Illumina libraries were prepared [16] and 125bp PE reads were generated on the Illumina HiSeq v4.

<u>Deep sequencing</u> was applied to quantify parasite load in batch 1 of Bone Marrow (BM) samples. Genomic DNA was sheared into 150bp fragments by focused ultrasonication (Covaris Inc., Woburn, USA) and libraries were prepared using the SureSelect Automated Library Prep Kit (Agilent, Santa Clara, USA). Index tagged samples were amplified using KAPA HiFi DNA polymerase, (KAPA Biosystems, Wilmington, USA) and quantified using an Accuclear DNA Kit (Biotium, Fremont, USA), 75bp PE reads were generated on the Illumina HiSeq 2500.

<u>SureSelect</u> was used for *Leishmania* genome capture. Genomic DNA was sheared into 150bp fragments by focused ultrasonication (Covaris Inc., Woburn, USA) and libraries were prepared, either as described above, or using the NEBNext Ultra II DNA Library prep Kit (New England Biolabs, Ipswich, USA). Samples were tagged with a unique index, amplified and quantified by qPCR (S1 Text). Samples were then pooled into batches for SureSelect enrichment, based on qPCR estimates of parasite load, so that each batch had as little variation in parasite load as possible: we aimed for only a 2-fold variation within a batch, while optimising the use of the sequencing data. Based on our experience of enrichment levels obtained in the experimental samples, batch sizes varied from 6 or 7 samples per batch where they had less than 0.05% *Leishmania* gDNA content up to 8 samples for higher parasite load. Samples were pooled within batches in an equimolar fashion. Pooled material was taken forward for hybridization, capture and enrichment using the standard SureSelect Target Enrichment System XT2 protocol recommended by the manufacturer (Agilent Technologies https://www.agilent.com/cs/library/usermanuals/Public/G9630-90000.pdf). Custom designed oligonucleotide baits were used at a 1:10 stock dilution. 75bp paired-end reads were generated on the Illumina HiSeq 2500.

<u>Amplicon sequencing</u> was used to explore the presence of minor genetic variants in clinical samples. In the first step we used the primers listed in Text S1 to amplify fragments that contained diagnostic SNPs/insertion for ISC3, ISC4, ISC5 and ISC5 groups using Advantage 2 Polymerase Mix (TAKARA, Kusatsu, Japan). 2µl of template with varying DNA concentration were used in a total of 50µl PCR reaction (S1 Dataset A). Touch down PCR with the following conditions was used: 95°C for 5 mins followed by 16 cycles of 95°C for 30s, 68°C to 6O°C (decrease of 0.5°C with each cycle) for 30s, 68°C for 30sec, followed by 19 cycles of 95°C for 30s, 60°C for 30s, 68°C for 30s and a final step at 68°C for 10min. Then, a second round amplification was performed, using 5µl of template, to add indexed Illumina adapters [17]. PCR conditions of the second round were 95°C for 3 mins followed by 10 rounds of 98°C for 20s, 65°C for 30s, 72°C for 30s finishing with 5 mins at 72°C. The final products were purified using AMPure XP SPRI beads and quantified on a plate reader using an AccuClear DNA quantification kit (Biotium, Fremont, USA) followed by equimolar pooling. Due to low base diversity of the amplicons, 20% PhiX was spiked into the resulting pool and 300bp PE reads were generated on the Illumina MiSeq.

### DNA read mapping, SNP and indel calling

SuSL-seq reads from clinical samples and data from 204 (54 strains, 150 isolates) previously sequenced *L. donovani* genomes [8] were mapped to the improved reference *L. donovani* genome LdBPKv2 [11], using Smalt v7.4 (http://www.sanger.ac.uk/science/tools/smalt-0). The methods were reported elsewhere [11]. Briefly, Smalt options for exhaustive searching for optimal alignments (-x) and random mapping of multiple hit reads were used. Reads were mapped when mapping base identity was greater than or equal to 80% to the reference genome (-y 0.8), to eliminate spuriously mapped reads mainly from human DNA. For the sequence analyses, all reads mapped to the reference, not limited to bait locations, were analyzed. For phylogenetic analyses and for evaluating the accuracy of SuSL-seq data, we used the population SNP and indel calling mode of UnifiedGenotyper in Genome Analysis Toolkit v3.4 with the SNP cut off 4000 used (GATK: https://software.broadinstitute.org/gatk/). No lower or higher depth cut off was used in the population SNP calling since the depth of some SuSL-seq samples was quite low.

### Reference genome masking and SNP screening

To mask repetitive regions and regions prone to generate false positive SNPs due to the presence of non-*Leishmania* DNA in clinical samples, we masked 7,453 regions that spanned a total of 43,200 bases. These regions were identified by blast similarity hits [8] and by using SNPs that were generated by pure human DNA SureSelect analysis and samples with lower amount of *Leishmania* DNA. The masked positions were provided in MaskSuSL4.bed file. We also excluded a SNP cluster containing more than 7 SNPs within 60 neighboring bases to exclude false SNPs from simulated *Leishmania* DNA samples mixed with mouse DNA Mpd1 and Mpd2. This was effective to eliminate non-*Leishmania* reads with a dozen of SNPs but with highest mapping score 60.

### Somy estimation

For somy estimation, 2 different methods were used. For samples with consistently high-quality mapping against *Leishmania*, such as genomic libraries from cultured promastigotes we used the median depth across each chromosome [11, 18]. For samples whose read coverage depth was more variable, such as amplified and unamplified clinical samples, depth was calculated in 1000bp windows to allow a more uniform analysis of coverage variation [11]. Previous studies have shown some correlation between read coverage depth and chromosome length for some sequencing runs [11]. To correct this bias, we applied local median correction [11]. This method calculates somy based on the median coverage values of chromosomes with similar lengths. In the calculation of somy, the first 7,000 bases and last 2,000 bases of each chromosome were excluded because these regions tend to be repetitive telomeric regions and their depth was not reliable. Similarly, certain over-amplified chromosome regions were excluded from depth calculation for SuSL-seq samples: these regions were listed in S1 Dataset B. The range of monosomy, disomy, trisomy, tetrasomy, and pentasomy was defined to be the full cell-normalized chromosome depth or S-value: S < 1.5, 1.5 ≤ S < 2.5, 2.5 ≤ S < 3.5, 3.5 ≤ S < 4.5, and 4.5 ≤ S < 5.5, respectively [11].

### Gene copy number variation

Local copy number variants were detected as described elsewhere [8]. In brief, we calculated an average normalised haploid depth of H-locus (Ld23: 90426 - 104470) and M-locus (Ld36: 2558460 - 2576450) for each strain and then obtained an average normalised haploid depth and standard deviation for each group. For ISC1-specific CNVs, we calculated an average normalised haploid depth for each gene for each ISC1 strain and then obtained an average normalised haploid depth and standard deviation for each CG group, using the depth of 191 sequenced strains (promastigotes). ISC1 specific gene CNVs were then defined to be genes whose normalized haploid depth differed more than ± 0.3 from the baseline average depth of other ISC groups. The value 0.3 was chosen here because we have observed many long CNVs in trisomic chromosomes, which were amplified or deleted in 1 of the 3 homologous chromosomes.

### Phylogenetic analysis

For phylogenetic analysis, 197 variables sites represented by 394 bases were used to perform neighbor-joining analysis using the p-distance model with 1000 bootstrap replicates as performed in MEGA 7.0.21 [19] and and Splitstree v4 [20]. These analyses used 191 promastigote strains plus 51 clinical samples (one clinical sample from ISC1 was excluded because phylogenetically too divergent). Artificial mixtures of *Leishmania* and human DNA (Leishmania DNA percentage of 0.06%) were used to measure the accuracy to multiple infections of the phylogenetic analyses.

### Identifying minor ISC genotypes and polyclonal infections with SuSL-seq reads

Two methods were used to identify minor ISC genotypes, respectively based on (i) SNP motifs specific of ISC groups and (ii) alternative allele frequency from sequenced reads.

We first extracted genotype specific 23-mer motifs that contained a SNP or reference base at the center. For ISC3, ISC4, ISC5, ISC6 and ISC7 groups, we created 26, 53, 38, 26 and 14 motifs that matched SNPs common to these groups, respectively. These group specific SNPs were identified using 191 samples analyzed in the previous study [8] and mapped to the pacbio reference [11]. To identify minor genotype reads which may have been missed by the standard SNP calling, we counted the reads in the alignment files that contained genotype specific motifs. These motifs were listed in S1 Dataset A and used to estimate the proportion of ISC3, ISC4, ISC5, ISC6 and ISC7 in each sample.

In the second approach, a GATK population VCF file for bone marrow and promastigotes were converted to a table that contained alternative allele frequency. In addition, total read depth, reference read depth, alternative allele read depth and genotype information created for VCF file were kept along alternative allele frequency. We checked alternative allele frequencies at group specific SNP sites to identify polyclonal samples. A mean of alternative allele frequency was calculated by averaging alternative allele frequencies at all ISC specific SNP sites per sampling, excluding missing depth sites.

### Identifying minor ISC groups using Amplicon sequencing

We applied a read count method and an exact motif matching method to identify minor ISC groups using amplicon sequencing. The read count method was based on samtools pileup and included reads marked as PCR duplicates, counting all four bases to access back ground base error rates. The exact motif matching method was based on a 17 mer for each diagnostic SNP and a 22 mer for the diagnostic insertion and they were listed in S1 Dataset C & D.

### Parasite cultures and competition experiment

Twelve clinical isolates paired to bone marrow samples were cultivated on HOMEM. Three *L. donovani* cloned strains were used in this experiment BPK067/0/cl2, BPK077/0/cl5, BPK091/0/cl9, respectively belonging to the ISC3, ISC6, and ISC4 genetic groups [8]. The parasites were grown on Tobie’s blood agar [21] with a saline overlay at 26°C to mimic the growth conditions during parasite isolation from clinical samples [22]. The parasites were inoculated to the concentration of 5×10^5^ml^-1^, and passaged to fresh medium every 3-4 days. For each strain mixture, 3 replicates were prepared, and the pure strains were grown in duplicates. The proportion between two parasites strains was roughly verified at the start of the experiment, and was further assessed approximately every 5 passages.

To follow the proportions between two different parasite strains in culture, a diagnostic SNPs, which allow distinction among different ISC genetic groups were used [23]. Specifically, the Ld36_p5263_F1* and Ld36_p5263_R1, and Ld33_p3644_F1 and Ld33_p3644_R1* primer pairs were used to distinguish between ISC3 and reference, and ISC6 and reference genotypes, respectively. The presence of these alleles was verified by standard Sanger sequencing, where primers mentioned above with * were used to sequence the amplified products. PCR amplicons were sequenced at Baseclear (Leiden, The Netherlands) with ABI 3730 (XL) DNA Analyzer, and the chromatograms were analyzed using the 4peaks software (Nucleobytes). The ability to detect both alleles was first verified by sequencing the PCR products amplified from DNA mixed in the same proportions as the parasites in the competition experiment. With this method, it was not possible to detect the exact proportions of the two alleles, but the relations between them (i.e. which allele was present in higher proportion) was assessed.

## Results

### SuSL-seq analysis of clinical samples: sequencing statistics

Details on the SureSelect bait design are described in S1 Text and in S1 Fig. A. SuSL-seq was optimized on artificial mixtures (*Leishmania* DNA diluted in mammalian DNA in different ratios: see details and metrics in S1 Text and S1 Fig. B-C and S2 Fig.): a percentage of *Leishmania* DNA of 0.006% was found to be the lowest limit for suitable analysis of genome diversity. SuSL-seq was then applied to 63 clinical samples with *Leishmania* DNA percentage of at least 0.006%, 59 bone marrow and 4 splenic aspirate samples collected from VL patients in Nepal (S1 Table) in the same region where we previously studied genome diversity of *L. donovani* isolates (S3 Fig.).

Among all SuSL-seq samples, the median depth of reads mapped against the *L. donovani* reference genome was 22.5, the 5^th^ and 95^th^ percentiles of median depth were 2 and 74.16, respectively, while the maximum depth was 126.5 (in BPK89BM) and the minimum average depth was 0.074 (in BPK161BM). Across all clinical SureSelect libraries, the mean percentage of reference genome bases covered by at least 5 reads was 61.1% and, the mean percentage of bases covered by at least 1 read was 78.4%. This coverage average was skewed by a small number of lower-quality samples. For instance, within a sequencing batch of 24 samples, over 90% of the genome was covered by at least one read in 14 samples; in all cases better genome representation was obtained following SureSelect enrichment (S4 Fig. A). Overall, as for the artificial mixtures, we obtained the highest enrichment for samples with lowest parasite loads. An approximately linear relationship was found between Leishmania DNA percentage in clinical samples and the proportion of reads mapping to *Leishmania* after SuSL-seq for samples with up to 1 % of *Leishmania* DNA (Fig. 1A). For samples with higher *Leishmania* content, no further increase was observed beyond around 60-70% of reads aligning to BPK282v2 reference genome (Fig. 1A). Eleven samples had low coverage, with less than 10% of the genome covered at a depth of 5x (Fig. 1B). These low-coverage samples also had the lowest parasite load, and were excluded from the analyses described below, so 52 samples were studied in greater detail.

**Fig. 1.**
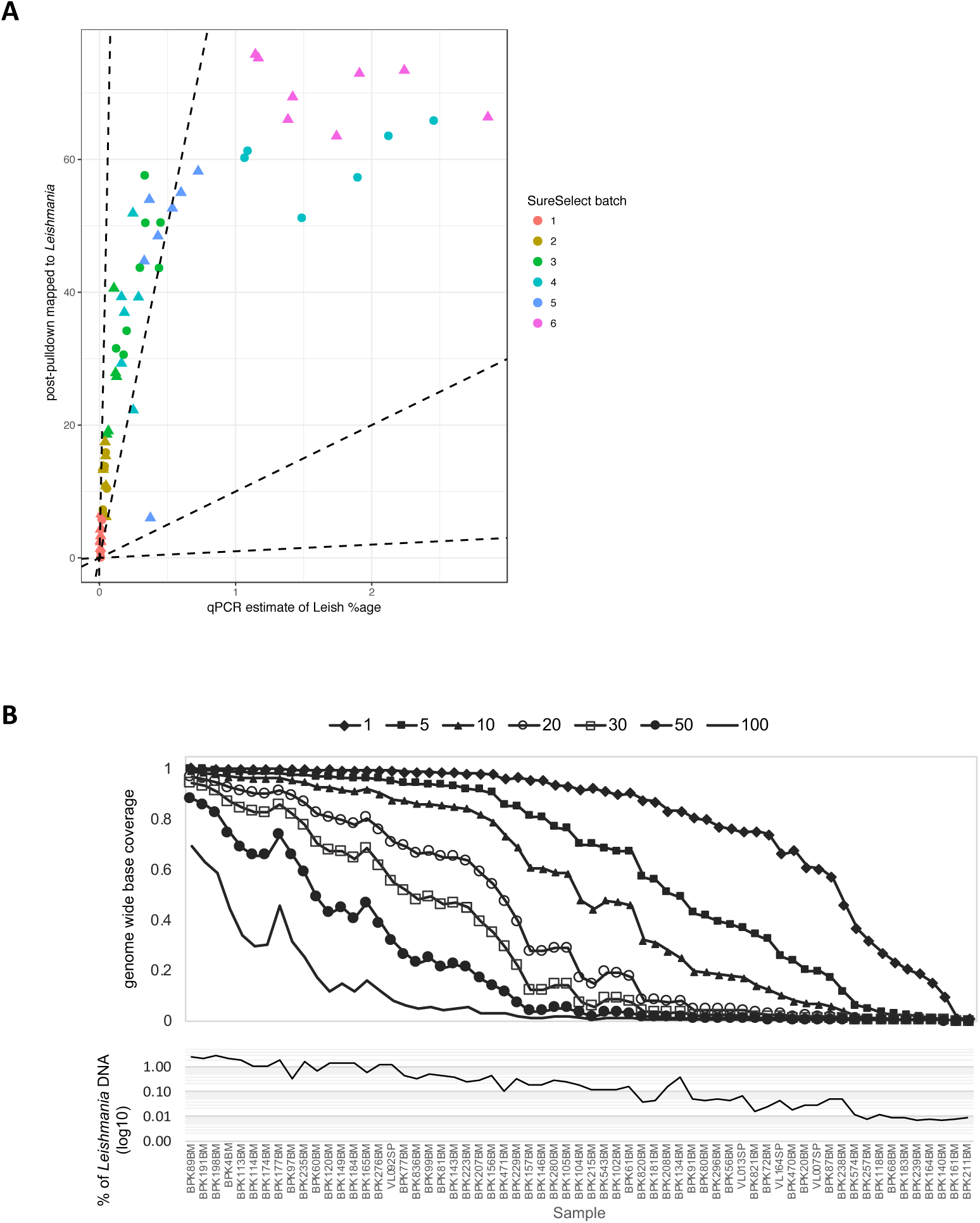
Performance of SuSL-sequencing on clinical samples. **A.** Enrichment of *Leishmania* DNA from clinical samples by SureSelect; The y-axis shows percentage of reads mapping to the *L. donovani* reference genome following SureSelect enrichment. The x-axis is the pre-enrichment percentage of *Leishmania* DNA, estimated by either whole-genome sequencing or qPCR; dotted lines indicate no enrichment and 10-, 100- and 1000-fold enrichment. **B.** Read mapping statistics in the 63 samples after SureSelect enrichment. Upper panel shows evenness of genome coverage as the proportion of bases covered with a minimum of number of sequencing reads from 1 to 100 as indicated by the seven different lines. Lower panel shows the percentage of reads mapping to the *Leishmania* reference genome for each sample, shown on the x-axis. The x-axis is common to both panels. BM, Bone Marrow; SP, Spleen aspirate.

### SuSL-seq analysis of clinical samples: genetic variants

Using an improved version of the BPK282 reference genome BPK282v2 [11], we were able to call SNPs in these 52 samples, of which 51 were identified as belonging to CG and one belonging to the genetically distinct ISC1 sub-population [8]. We identified 197 variable sites that had sequence coverage and could therefore be reliably genotyped in all 51 CG clinical samples; these sites were also characterized in 191 previously sequenced parasite isolates from the CG [8] and 9 newly sequenced isolates (see below). The 197 sites were sufficient to reconstruct a phylogeny consistent with our previously published analysis [8], placing all clinical SureSelect samples within the previously identified ISC diversity for *L. donovani* (dots in Fig. 2A). Out of the 51 CG samples, 40 clustered in one of the previously defined groups (ISC3-6, and ISC9). Five samples clustered at the base of ISC4. Six clinical samples, clustered with isolates previously designated as ‘ungrouped’ (Fig.2A and S5 Fig. A-B), forming a novel monophyletic group that we have named ISC11. We verified the accuracy of our experimental and bio-informatic pipeline by including in the phylogenomic analyses SuSL-seq data from artificial mixtures (DNA of BPK282 diluted in human DNA): as expected, they clustered in the ISC6 group with the original BPK282 sequence data (not shown).

**Fig. 2.**
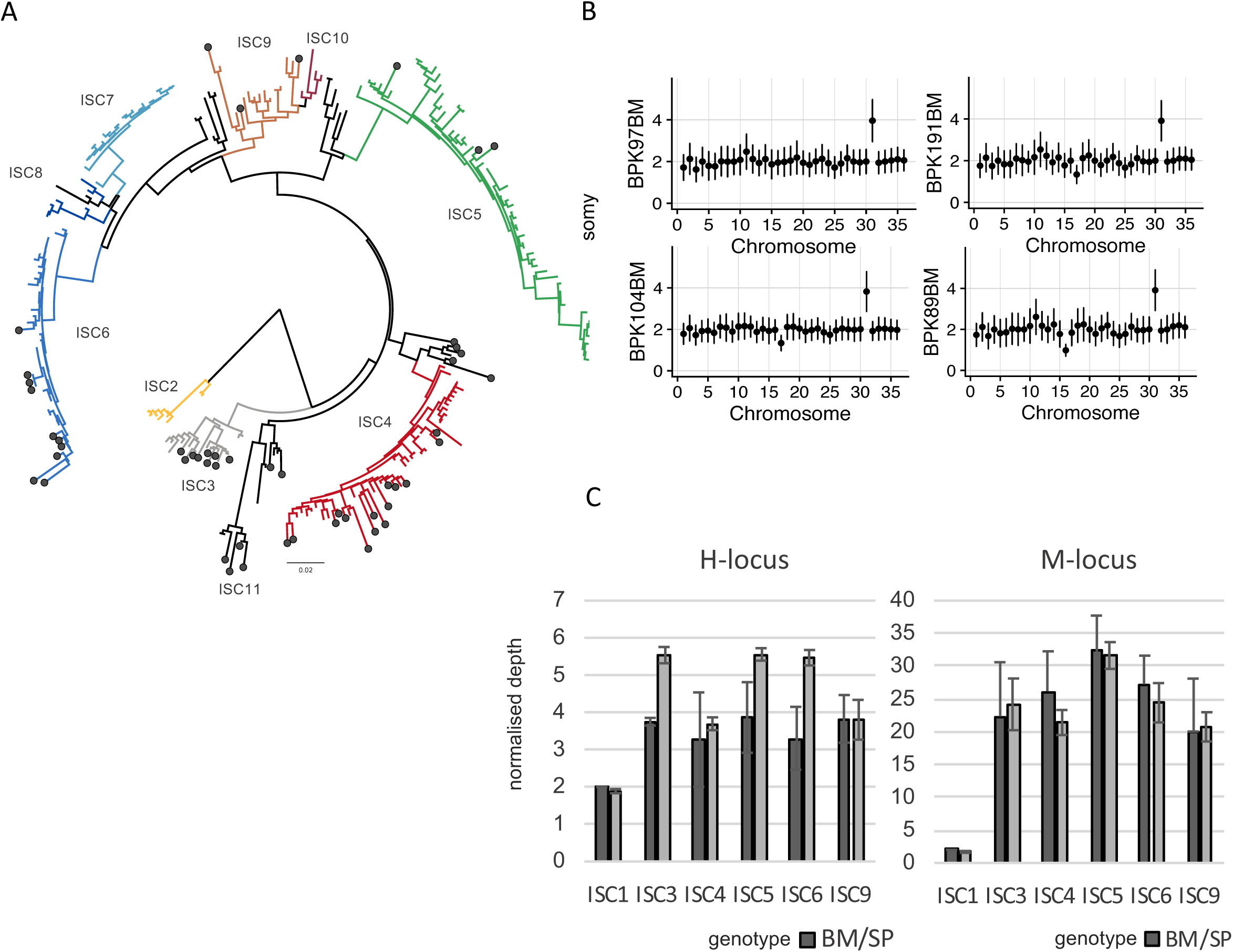
Genomic diversity among clinical samples. **A.** SNPs. Phylogenetic tree based on SureSelect enriched clinical samples (black circle) and previously sequenced isolates (1). Diagram is a neighbor-joining phylogeny based on 197 variable sites. ISC2-ISC10 were sub-populations previously defined (1); ISC11 is a novel group that includes clinical samples and isolates previously designated as ‘ungrouped’. Full sample identifiers are shown in Fig. S5A. **B.** Karyotypes (High quality samples). Y axis shows normalized somy estimate for each chromosome (x axis). Points show central estimate and bars show one standard deviation around these estimates, calculated by the binned depth method. **C.** Local CNVs. Average copy number per cell of H- and M-loci per ISC group in SureSelect enriched clinical samples (Bone Marrow or Spleen aspirate; BM/SP) and in cultured isolates (promastigotes, Prom). Error bars show one standard deviation around the mean estimate. Only a single sample was available from bone marrow aspirates in ISC1, hence no standard deviation is shown.

Notably, among the 11 samples that were excluded from the phylogenomic analysis because of poor genome-wide coverage, nine could be attributed to a specific ISC group, on the basis of diagnostic SNPs specific to these groups (labeled with * in S1 Table). The remaining two samples, BPK211BM and BPK161BM, had sequence data with too low coverage for any genotyping (S1 Dataset E). Analysis of genotype distribution per village revealed sympatric genotypic diversity: for instance, in Itahari, four different ISC types were sampled over 11 days in June 2002 and three were sampled over four months in 2003 (S1 Dataset F). Interestingly, in this village, two cohabiting members of the same family were sampled during each of these two periods: in both cases, two different ISC types were identified. We also searched for small indels that were unique to clinical samples. We identified six unique indels: five were located in non-coding regions, and one introduced a 1-bp frameshift in the LdAQP1 gene (S1 Dataset G).

Next, we examined chromosome copy numbers. Consistent with our observations in artificially mixed samples (S1 Text), the apparent accuracy of our somy estimates in clinical samples depended on the *Leishmania* DNA percentages, as those with low percentage had lower normalized read depth and genome coverage (S1 Dataset E). For example, chromosome 31 is known to be tetrasomic in all *Leishmania* species and isolates sequenced so far, except under experimental drug resistance selection where somy can be even higher [9]. As expected, S-values (normalized coverage depth, see Methods) of chromosome 31 were close to 4 in samples with high genome coverage. These values, however, were lower in samples with lower genome coverage, even though the somy of chromosome 31 was distinctively higher than 2 except for the very lowest coverage samples. To distinguish real somy variation and technical artefacts, we classified samples into group of similar genome coverage into those with high (36 samples, genome coverage above 89.1%), medium (16 samples, coverage 56.6-86.9%) and low quality (9 samples, coverage 9.8%-45.8%) somy calls (S1 Dataset A and S6 Fig. A). Two samples (coverage 1.1% and 0.8%), were excluded from this analysis because their depth was too low to quantify somy. The high-quality samples provided a clear picture of the average ploidy of amastigote populations in human hosts: most chromosomes showed an S-value around 2 (thus overall disomic), with only chromosome 31 showing consistently high somy around 4 (Fig. 2B). Other exceptions were chromosomes 16 and 18 which showed S-value of 1 (thus overall monosomic) in BPK89BM, and in BPK146BM, respectively (Fig. 2B and S6 Fig. A1). Chromosome 17 showed a S-value of 1 or lower than 2 (suggestive of mosaicism between monosomic and disomic variants) in seven samples: BPK276BM, BPK104BM, BPK146BM, BPK161BM, BPK198BM, BPK191BM, BPK4BM (Fig. 2B and S6 Fig. A1-3). We detected S-values between 2 and 3 (potentially indicating the presence of mosaicism between disomic and trisomic variants) of chromosome 20, 23, 32 and 33 in BPK821BM. (S6 Fig. A1).

We detected two main local copy number variants (CNVs) specific of CG (termed M- and H-locus, [8]) in all 51 SuSL-seq CG samples. When we compared average normalized depth for these two CNVs between clinical and previously published promastigote samples for each genetic group, we found good agreement for M-locus with correlation R^2^=0.941 and p-value=2.96 x 10^-4^, and slightly lower depth for H-locus in clinical samples with correlation R^2^=0.571, p-value=4.84 x 10^-2^ (Fig. 2C). The ISC1 group contained many other CNVs that were used to test the accuracy of local depth measured in SureSelect platform. We found correlation (R^2^=0.9921, p-value=5.3449e-14) between the depth in the ISC1-specific CNVs in the previously analyzed promastigote samples and the depth in the BPK72BM clinical sample (S6 Fig. B).

### Comparison of paired clinical and cultured isolate samples

For 12 of the SuSL-seq clinical samples, paired cultured parasite isolates derived from the same clinical material were available. We compared the genotypes within each of these 12 pairs, at three levels: SNPs, indels and chromosome copy number. SNP analysis revealed that nine SuSL-seq samples were assigned to different ISC genotype groups to the matched cultured isolates (Table 1), with samples assigned to different core group genotypes differing by 9-26 homozygous sites and 3-13 heterozygous positions. In two cases, the clinical samples were genotyped as being in the CG and the isolate was assigned to ISC1; these pairs of samples were far more distinct, differing in 29942 and 34146 homozygous SNPs and 4370 and 167 heterozygous sites respectively. For the 3 paired samples with matching ISC genotypes, we found only minor nucleotide differences (0-9 homozygous and 3-7 heterozygous SNPs). We found differences in Indel variants in 10 out of 12 samples: in one of them (BPK081) a single base insertion was observed in LdAQP1 (described in detail above) in the bone marrow, which was not present in the paired culture (Table 1). Finally, chromosome copy number differed in 10 paired samples, while in BPK087 and BPK081 we found no significant differences in somy values, and parasites remained overall disomic in culture (Fig. 3A). Accordingly, none of the 12 paired samples revealed identical genomic features between the clinical samples and the derived isolates.

**Table 1.** Genotype differences observed in 12 paired samples: clinical samples analyzed by SureSelect and derived isolates sequenced classically. Total base difference* = Homozygous difference x 2 + Heterozygous difference. Almost diploid**: S-value was 2 in all chromosomes except the tetrasomic chromosome 31. Notes: (1) the S-values of chromosome 11 in all bone marrow samples were slightly higher due to amplification artefact and therefore, their values were assumed to be disomic. (2) The S-values of chromosome 31 of BPK87BM and BPK80BM were lower than the expected tetrasomic values due to lower genome coverage.

| Clinical sample ISC genotype | Derived isolate ISC genotype | Total base difference* | Homozygous difference | Heterozygous difference | Indel difference | Karyotype (bone marrow) | Karyotype (derived isolate) |
| --- | --- | --- | --- | --- | --- | --- | --- |
| ISC4 | ISC6 | 47 | 20 | 7 | 4 | almost diploid** | trisomy on ch5 and 11 |
| UG | ISC4 | 42 | 17 | 8 | 3 | almost diploid | trisomic on ch2, 4, 7, 8, 9, 11, 12, 20, 26, 33 and 35. tetrasomic on ch23 |
| ISC6 | ISC4 | 55 | 21 | 13 | 4 | almost diploid | regular |
| ISC3 | ISC4 | 36 | 14 | 8 | 8 | almost diploid | trisomic on ch20, 26 and 33 |
| ISC3 | ISC1 | 68459 | 34146 | 167 | 6275 | almost diploid | monosomic on ch17 |
| ISC4 | ISC1# | 64254 | 29942 | 4370 | 8281 | almost diploid | trisomic on ch23, 33 and 35 |
| ISC9 | ISC5 | 55 | 26 | 3 | 3 | almost diploid | trisomic on ch2, 5, 8, 9, 11, 12, 13, 14, 20, 23, 27 and 32. tetrasomic on ch33 |
| ISC5 | ISC4 | 40 | 18 | 4 | 1 | monosomic on ch17 | trisomic on ch4, 11, 20, 23. tetrasomic on ch5 |
| ISC4 | UG | 21 | 9 | 3 | 4 | almost diploid | trisomic on ch1, 7, 8, 10, 13, 15, 20, 22, 23. tetrasomic on ch9 |
| ISC4 | ISC4 | 13 | 3 | 7 | LdAQP1: frameshift G/GA insertion | almost diploid | regular |
| ISC6 | ISC6 | 5 | 0 | 5 | 0 | almost diploid | trisomic on ch8, 9, 13 and 23 |
| ISC4 | ISC4 | 11 | 1 | 9 | 0 | almost diploid | trisomic on ch1, 7, 8, 10, 13, 15, 20, 22 and 23. tetrasomic on ch9 |

**Fig. 3.**
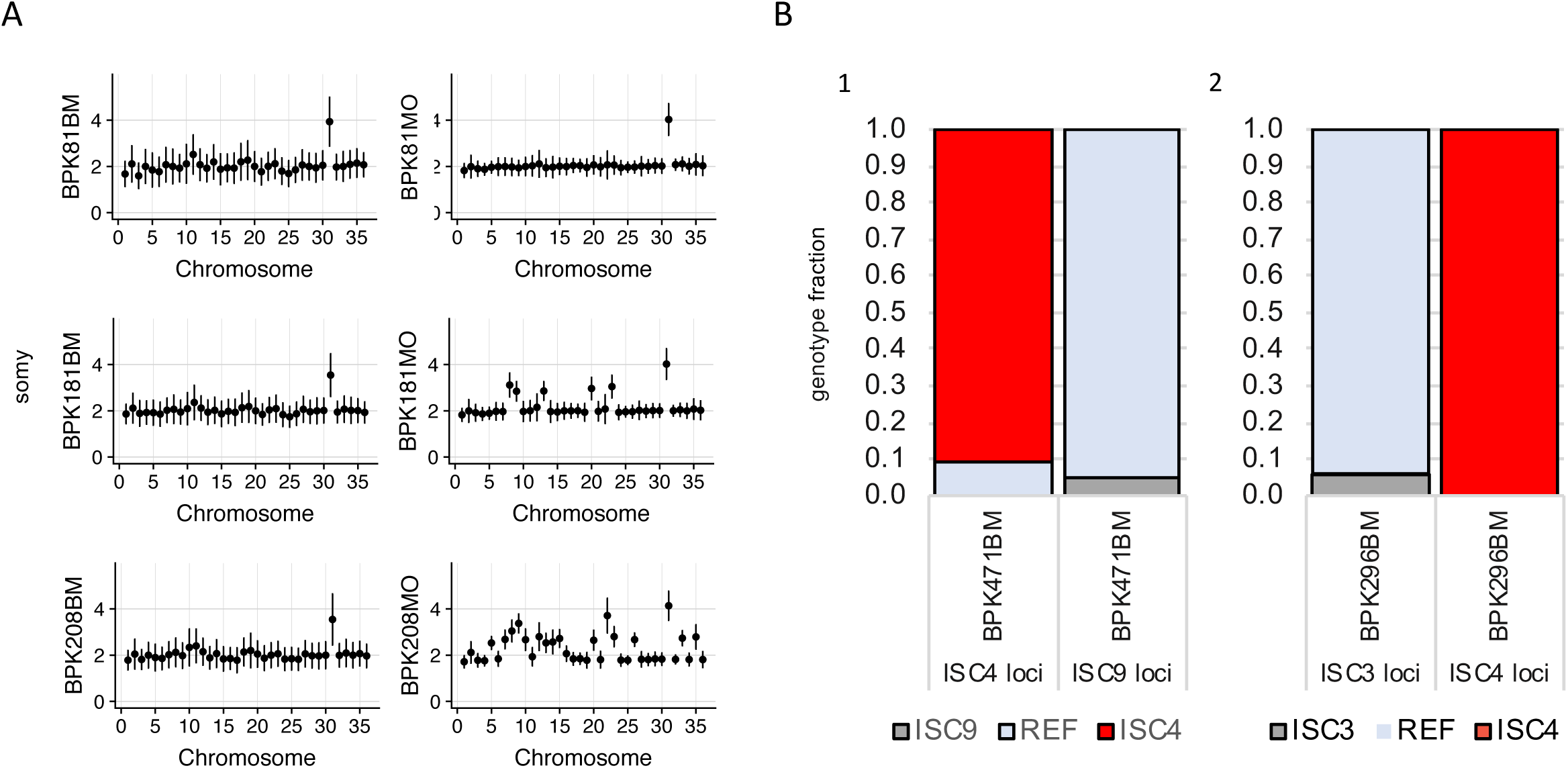
Somy comparison between matched clinical samples and culture parasites. **A.** Inferred somy for 3 samples for which SureSelect-enriched bone marrow (BM) samples and cultured isolates (MO) from the same patients had matching ISC genotypes. The y-axis shows normalized somy estimate for each chromosome (the x-axis). Points show central estimate and bars show one standard deviation around these estimates. Somy estimates and standard deviations for the bone marrow samples were based on the binned depth method while corresponding values for solates were based on depth of each position. **B.** Evidence for polyclonal infections. The bars represent the proportion of sequencing reads showing the ISC-specific genotype or the reference (REF) genotype at loci with ISC-specific alleles. 1) Genotypes at ISC9 and ISC4 loci for SureSelect-enriched bone marrow sample BPK471BM. 2) Genotypes at ISC4 and ISC3 loci for SureSelect-enriched bone marrow sample BPK296BM.

We then focused on the nine paired samples for which different ISC types were detected in the bone marrow and the derived isolates. Considering the sympatric ISC genotype diversity highlighted above, it is possible that some patients harbor polyclonal infections, with a given genotype dominant in a human host and another one dominant in culture, because of fitness differences in the respective environments. This possibility was explored by further genetic analyses and by competition experiments.

First, we found a second and minor genotype in the SuSL-seq mapped reads of two clinical samples (BPK471BM and BPK296BM), at a proportion estimated as 9.2%, and 6%, respectively (Fig.3B). We also found the evidence of polyclonality in one of the cultured isolates, which differed in its ISC genotype from the paired clinical sample (Table 1): in BPK157MO, ISC1 constituted the main genotype (93.85% of the reads), and we found 6.15% that were not ISC1 at all ISC1 specific SNP sites. Secondly, we tested for the presence of more than one genotype in un-enriched clinical samples by applying ISC single locus genotyping (ISC-SLG) combined with NGS-based deep amplicon sequencing using four previously described diagnostic SNPs/insertion allowing discrimination between the ISC3, ISC4, ISC5 and ISC6 genotypes [23]. Short regions flanking the SNP/indel were amplified, sequenced and the frequency of both alleles in all the tested samples was estimated. In order to increase the chance of detecting the presence of minor genotypes, we selected clinical samples with the highest parasite loads and/or highest total DNA contents. In addition, we included polyclonal clinical samples BPK471BM and BPK296BM (see above), two clinical sample-isolate pairs (BPK077 and BPK276) and cloned lines, which served as positive or negative controls of the presence of polymorphism at the diagnostic SNP/indel sites. The estimated percentage of *Leishmania* DNA and number of *Leishmania* genome equivalents in the clinical samples varied between 0.05-2.85%, and 12-60901, respectively (S1 Dataset A). We could confirm the principal ISC genotypes defined by SuSL-seq in all clinical samples (S1 Dataset A). Alternative alleles containing SNP variants could be observed in most of the clinical samples, but their frequency was not significantly different from the frequency found in the control cloned lines (S7 Fig. and S1 Dataset A). For the AQP1 insertion, the situation was similar with the exception of the sample BPK120BM (defined as belonging to ISC9), which indicated the presence of a small population of ISC5 genotype (z-score:3.2; p-value: 0.0014). The reciprocal was found in BPK276BM, an ISC5 sample in which a small proportion of WT alleles was encountered and we found no alleles containing the ISC5-specific AQP1 insertion in the paired clinical isolate. Although these results are not significant (p-value: 0.139), they are consistent with the observed ISC5 genotype in BPK276BM, and ISC6 genotype in the derived isolate (S1 Dataset A).

Finally, to test whether different growth rates could explain the different genotypes observed between direct clinical samples and those grown in vitro, we set up a growth-competition experiment. We inoculated flasks with cloned strains BPK091 and BPK077 (ISC4 and ISC6, respectively), and BPK067 and BPK091 (ISC3, and ISC4, respectively) together in the following ratios (strain1/strain2): 90/10, 50/50, and 10/90. Using Sanger sequencing to monitor the presence of diagnostic SNPs that could distinguish the two genetic groups at different time points during *in vitro* cultivation, we were able to detect which of the two co-cultured strains was dominant. We reproducibly observed for both parasite combinations that one strain consistently dominated the culture, even if it was only present as 10% of the initial inoculum (S8 Fig.). Interestingly, BPK091, which was present in both mixes, became dominant after 21 passages when co-cultured with BPK077 but after only five passages it was out-competed by BPK067.

## Discussion

We demonstrated here proof-of-principle for the sequencing of *Leishmania* genomes directly from clinical samples of patients with VL. Tested on 63 clinical samples from Nepal, SuSL-seq showed a high analytical sensitivity: we were able to (i) perform phylogenomic analyses on 82.5% of samples, (ii) assign 97% of samples to previously defined ISC genotypes using a set of diagnostic SNPs, (iii) estimate chromosome copy number in 82.5% of samples and (iv) identify large local CNVs in 83%. With the current design of the baits, our method should be applicable to all the parasites causing visceral leishmaniasis i.e. *L. infantum* and *L. donovani*, in East Africa, the Mediterranean basin and Latin America. Further work is required to test SuSL-seq on other host samples (animal reservoir, insect vectors, or different host tissues). It would be desirable to improve its performance in *Leishmania* samples with low parasite load and/or low amount of total DNA: here, the combination of SureSelect with other enrichment methods might prove useful.

Our study sheds new light on the biology of *Leishmania* in the human host. A first aspect concerned the karyotype of amastigotes. Massive aneuploidy has been described in cultivated *Leishmania* promastigotes of all species [24]. In particular, sequencing of 204 *L. donovani* genomes [8] revealed strong differences in karyotype between parasite strains, with a baseline overall disomy and up to 22 polysomic chromosomes (essentially 3N and 4N) in single strains. In sharp contrast, we did not find high levels of aneuploidy in the amastigotes present in clinical samples of the same region. Besides chromosome 31 which is constitutively tetrasomic in all *Leishmania* species, most chromosomes in amastigotes of clinical samples were inferred to be disomic, and only a few monosomic or with intermediate coverage depth, the latter likely reflecting mosaic aneuploidy in the analyzed parasite populations, with mixtures of cells of different karyotypes [25]. Overall, the differences in aneuploidy observed between clinical samples and derived isolates are consistent with our previous findings where we reported close to diploid karyotype in parasites adapted to Syrian golden hamster, and highly dynamic aneuploidy during propagation of *L. donovani* in vitro [11]. This genomic plasticity was mirrored at transcriptome level and might provide a rapid adaptation in response to environmental cues.

A second potential finding is related to the observed genotypic differences between 12 clinical samples and derived cultured isolates. Strictly speaking and considering different types of genetic variants (SNPs, indels an somy), none of the 12 isolates showed an identical genotype to the parasites present in the paired bone marrow samples. We cannot exclude that the 1-nt indel in the LdAQP1 of BPK81BM ‘mutated’ during culture maintenance and restored the reading frame in the corresponding isolate’s genome. However, knowing that frame shifts in this gene have been linked to *Leishmania* antimony resistance [8], this result shows the risk of losing drug resistance markers when genotyping isolates. The differences in karyotype as observed in 11/12 paired samples could also be acquired during culture, but could also have resulted from selection from a mosaic background, as shown experimentally [25]. More generally, however, the SNP differences that led to isolates and SuSL-seq samples representing different ISC genotypes did not appear to be explained by mutations appearing *in vitro*; each ISC group is characterized by a set of diagnostic SNPs and the observed switch of ISC genotypes concerned all diagnostic SNPs in these sample pairs. In two cases, the isolate and clinical samples differed at tens of thousands of sites, and clearly represented the two very different populations of parasites circulating in VL patients in Nepal. We excluded SNP-calling artefacts, and the clinical work followed specific procedures for labeling, processing and tracking all samples. Thus, excluding experimental mistakes as the reason for these results, the most likely explanation is polyclonality of the infection in the host. In such a scenario, different clones would be most abundant in the host and the promastigote culture medium, due to their differences in respective fitness in these two environments.

We have attempted to verify the polyclonality hypothesis through a set of direct and indirect experiments. On one hand, in two clinical samples and one cultured isolate we detected minor reads (above 6%) with the signature of a second ISC genotype. We also tried to detect the second genotype directly in selected bone marrow samples with amplicon sequencing of 4 loci, which contain SNPs/AQP1 indel specific for different genetic groups from ISC. However, we were unable to find any statistically robust evidence for presence of a second genotype with this method, except in one case, BPK120BM. This is most likely due to a sensitivity issue of the amplicon sequencing method which was applied: (i) without enrichment (in most cases, in presence of more than 99% of human DNA, Dataset S1D), (ii) to find minor variants and (iii) targeting one single variant within (iv) a single copy gene. The two clinical samples BPK296BM and BPK471BM illustrate this problem well: they were clearly shown to be polyclonal from SuSL-seq data, while the second clone could not be detected from amplicon sequencing, most likely due to the low number of *Leishmania* genome equivalents (respectively, 43 and 12). On the other hand, the plausibility of this hypothesis was indirectly validated by a competition experiment in which we co-cultivated strains of different ISC genotypes in the same tube: even if a strain constituted only 10% of the initial inoculum, it could rapidly and in a reproducible way dominate over time in culture.

Further work is required to assess the extent of polyclonality among *L. donovani* infections. The phenomenon is common in *Plasmodium*, especially in high transmission areas [26]. Among trypanosomatids, it has been reported in *Trypanosoma cruzi*, where among others multiple genotypes were found to be transmitted congenitally [27] and temporal variation of genotypes was observed in asymptomatic carriers [28]. In *Leishmania*, mixed infections of different species were reported in wild rodents [29], horses [30] and humans [31, 32]. A recent study documented the possibility of polyclonality in *L. donovani* from East Africa, as different genotypes were isolated from different organs of the same patient [33]. A similar observation was made on a Spanish dog in which two different zymodemes of *L. infantum* were collected from the skin and a lymph node [34]. Furthermore, natural *Leishmania* hybrids are regularly reported all over the world (see for instance [35, 36]) and in our previous study in the ISC [8], we found 8 isolates of hybrid origin (out of 192 isolates of the CG, i.e. 4.2%). This could only occur in sand flies as a result of polyclonal infections.

The present work highlights the complexity of *L. donovani* infections in the human host. It appears that a patient can be infected by multiple clones showing different genotypes, which possibly vary at the somy level. This genetic diversity could offer an adaptive advantage to the whole population of cells, by providing ‘individual’ solutions to different environments, leading to different dominant genomes in the patient and in culture. We showed that this did not have a major impact so far for phylogenomic studies. However, if any link is to be made between parasite genome variation and clinical phenotypes – for example treatment outcome or pathogenicity – analyzing *Leishmania* directly from clinical samples is necessary, as both the genotype and aneuploidy in the mammalian host and in vitro culture can differ.

### Data Release

Raw data was deposited in the European Nucleotide Archive (ENA) with the accession number ERP110990

## Acknowledgements

This study was supported by the European Commission (EC-FP-222895), Belgian Science Policy Office (TRIT, P7/41), Flemish Fund for Scientific Research (G.0.B81.12), and Department of Economy, Science and Innovation in Flanders ITM-SOFIB (SINGLE project, to J.C.D., M.V.A. and M.A.D.). MB, MJS and JAC are supported by Wellcome via their core support for the Wellcome Sanger Institute (grant 206194). We acknowledge the support of the core pipeline staff at WSI, in particular Peter Ellis and Sara Widaa, and the help of Owen Hardy from Agilent technologies for assistance in designing the SureSelect array. The funders had no role in study design, data collection and analysis, or preparation of the manuscript.

## Supporting information

**S1 Text.** Supplementary results: SureSelect bait design, SureSelect genome capture and sequencing (SuSL-seq) optimization on artificial mixtures; Direct sequencing of *Leishmania donovani* genomes in 63 clinical samples. Supplementary methods: clinical samples, amplicon sequencing. Supplementary references. *(S1Text.docx)*

**S1 Table. List of Nepalese clinical samples studied here**: date of sampling and geographical origin, type of sample, treatment received (AmB, Amphotericin B; SAG, Sodium Antimony Gluconate; MIL, Miltefosine) and outcome, ISC group (according to the results of current study), *because of lower sequence quality, ISC genotype was defined based on a set of diagnostic SNPs, remarks about (i) treatment provided before admission or (ii) family links between patients, isolation outcome and code of successful paired isolates. *(available on request to authors)*

**S1 Fig. A. Distribution of probes across *L. donovani* BPK282 reference genome**. Most of the reference genome is covered by only a single probe. Some regions are not covered to avoid repetitive regions and regions with homology to mammalian hosts. Changes in the genome assembly subsequent to the bait design and the presence of additional probes to capture *L. infantum* have led to some regions being covered by multiple probes sequences. **B.** Much of the variation in depth in coverage in clinical samples is due to the presence of multiple probe sequences. Plots show the total coverage density across all clinical samples for regions of the reference genome covered by 0, 1, 2 or more probe sequences. **C.** Depth of read coverage in SureSelect enriched samples across two regions of the *Leishmania* genome. Lines show normalized coverage depth (actual coverage per base pair divided by genome-wide mean coverage) for one clinical sample (BPK543, blue) and the sum for all clinical samples (orange). Grey shading shows regions where SureSelect probes map to the genome. Mean coverage for BPK452 was 9.2 reads, for the summed clinical samples 2447.8. *(S1toS8figs.pdf)*

**S2 Fig. Performance of SuSL-sequencing on artificial mixtures of** *Leishmania* and mammal DNA at three different *Leishmania* DNA percentages: 0.06, 0.006 and 0.0006 %. **A**. Summary statistics for sequencing data from these experiments, showing the total number of reads in each library, the proportion of reads mapping to the *L. donovani* reference genome, and the enrichment factor, calculated as the ratio of the proportion of reads mapping to *L. donovani* in the SureSelect libraries to the proportion of *Leishmania* promastigote DNA included in the pre-pulldown DNA mixtures. **B.** Evenness of genome coverage. The y-axis shows proportion of bases covered with a minimum number of sequencing reads--read depth along the x-axis--by reads samples from SureSelect sequencing data from artificial mixtures. 104M, 78M, 52M, 26M and 13M indicate the total number of reads (in millions) sampled from each library; 0.06, 0.006 and 0.0006 are the input proportions of *Leishmania* DNA in percent. For lower *Leishmania* DNA percentages, the higher read depth did not result in higher genome base coverage. **C**. Inferred somy from SureSelect on artificial mixtures. BPK282, pure DNA from promastigote; LD06-0006, simulated clinical samples at three *Leishmania* DNA percentages (0.06%, 0.006% and 0.0006% respectively). Y axis shows normalized somy estimate for each chromosome (x axis). Points show central estimate and bars show one standard deviation around these estimates from four replicates of each *Leishmania* DNA percentages 0.06%, 0.006%, 0.0006%, respectively. *(S1toS8figs.pdf)*

**S3 Fig. Map of Nepal showing geographical origin of clinical samples and cultured promastigote isolates.** Colours represent different genotype groups defined elsewhere [8]. Circles represent Nepalese promastigote isolates from ref 8, crosses bone marrow and spleen samples. *(S1toS8figs.pdf)*

**S4 Fig. A. Genome sequencing of clinical samples: percentage of the genome covered by more than 1 read, with and without SuSL enrichment. B**. Relationship between qPCR estimate of *Leishmania* DNA concentration (y axis) and the proportion of sequencing reads mapping to the *L. donovani* reference genome before SureSelect enrichment in clinical samples for which both qPCR and pre-enrichment sequence data are available, confirming accuracy of the qPCR estimates. Batches were formed of samples with similar proportions of *Leishmania* DNA pre-enrichment. *(S1toS8figs.pdf)*

**S5 Fig. A. Phylogenetic tree (Neighbor-joining) based on bone marrow (BM), spleen (SP), isolates paired to clinical samples (MO) and previously sequenced lines (labelled A1 or B2, ref 8).** The tree is identical to the main one (Fig 2A), except that the labels of the samples are indicated. Bone marrow (BM) and spleen (SP) samples are labelled in blue; isolates that are paired to clinical samples are labelled in red, previously sequenced lines are labelled in black. ISC2-ISC10 were sub-populations previously defined (ref 8). See Fig.2A for comments on ISC11. **B.** Results from STRUCTURE analyses from five replicate runs (2×10^6^ MCMC chains following 10^6^ burn-in steps) under the Admixture model assuming 1-15 *K* clusters. The plot on the left shows the estimated loglikelihood of all runs for each *K* cluster. Barplots on the right summarize the assignment probabilities of every *L. donovani* sequence to each inferred cluster assuming 5, 6, 7, 8 or 9 clusters. Arrows denote the CG samples sequenced in this study. *(S1toS8figs.pdf)*

**S6 Fig. A. Somy in bone marrow and spleen samples.** BM=Bone marrow and SP=spleen. The y-axis and x-axis represent somy and chromosome, respectively. S-values and standard deviation error bars are given in black filled circle and vertical line, respectively. The S- and standard deviation values for the bone marrow samples were based on the binned depth method. The samples were sorted in the order of higher genome coverage, the 3 categories reflect resolution of somy estimation 1) High: whose genome coverage ranging from 99.45% to 89.10%. 2) Medium: whose genome coverage ranging from 86.93% to 56.60%. 3) Low: whose genome coverage ranging from 45.80% to 9.80%. In the lower Leishmania DNA concentration samples, it is still possible to identify higher copy number in chromosome 31. S-values of chromosome 2 and 11 were higher for almost all samples but these higher values were also observed in artificial mixtures of BPK282 where chromosome 2 and 11 were definitively disomic. Close inspection of read depth did not reveal any particular evidences to support their S-values to be higher than 2. **B.** Correlation between gene depth at 303 ISC1-specific CNV genes identified in promastigote samples. Correlation was moderately high r^2^ = 0.68, p-value = 10^-76^ and slope = 0.882 indicating that SuSL-seq can be used to confirm existing group-specific CNVs. On the plot, colored dots represent counts in log10 scale and probability density distributions were given on the corresponding axes. *(S1toS8figs.pdf)*

**S7 Fig. Allele frequency for the amplicon diagnostic of ISC5 genotype estimated with NGS-based deep amplicon sequencing.** Blue line represents the frequency for the non-ISC5 allele, while the orange line represents the ISC5-specific allele, which contains a 2bp-insertion in the AQP1 locus. Arrows indicate the two clinical samples in which a second allele was detected above the background, being statistically significant for BPK120_ISC5_BM. BM, Bone Marrow; SP, Spleen aspirate; MO, cultured isolate; NC, Negative Control, a cloned strain where the diagnostic ISC5-specific allele is absent; PC, Positive Control, a cloned strain characterized by the presence of the diagnostic ISC5-specific allele. *(S1toS8figs.pdf)*

**S8 Fig. Growth competition experiment.** Flasks were inoculated with individual cloned strains or mixtures of two cloned strains in the following ratios: 90/10, 50/50 and 10/90. Cultures were analyzed at the beginning of the experiment (Start), after 5, 15, 20 and 24 passages (P5, P15, P20 and P24 respectively). Diagnostic PCR that allows to distinguish between the two compared strains was used to monitor the presence of each strain in culture. A selected fragment of chromatogram containing the variable nucleotide (labelled with red circle) is shown. Arrows point at the samples were the changes in dominance occurred. **A**. Competition between the BPK091, and BPK077 strains (ISC4, and ISC6, respectively). **B**. Competition between the BPK091, and BPK067 strains (ISC4, and ISC3, respectively). *(S1toS8figs.pdf)*

**S1 Dataset. A**. ***Leishmania* genome equivalents in samples used for NGS-based amplicon sequencing.** The table shows the percentage of Leishmania DNA in the sample, the total amount of DNA, the amount of *Leishmania* DNA and the corresponding *Leishmania* genome equivalents used in each reaction to amplify short DNA fragments containing the diagnostic SNPs/AQP1 insertion for specific ISC genotypes. SuSL-seq genotype denotes the genotype assessed based on the phylogenomic clustering, while amplicon-seq genotype is based on the 4 diagnostic SNPs described by Rai et al. (ref 10). $, there are no ISC group-specific SNPs for ISC9 and ISC11, hence we could only report the negative results for ISC3-6 diagnostic SNPs. **B. Excluded regions for somy estimation**. For all chromosomes, the first 7000 bases and the last 2000 bases were excluded for depth evaluation. In addition, positions (x) in the following ranges were excluded because of the depth errors associated with SureSelect enrichment and because of the known major repetitive or amplified regions (see remarks). **C. Genotyping using group specific SNP motifs**: The table shows the ISC group, chromosome, position, reference allele, SNP allele, SNP score based on GATK, reference base motif and SNP base motif. All the diagnostic SNPs were homozygous SNPs except those of ISC7. Here ISC7 genotype was not detected: in a previous study (1), we found that genotype exclusively in India. A diagnostic SNP and reference base are located at the 12th base. **D. ISC5 AQP1 insertion motifs**: The reference base motif and AQP1 insertion base motif, along with neighboring motives that includes 2 base insertions of any base combinations at a given position are shown. The frequency counts of these motives were given for each sample and these neighboring motives were used to estimate the error rates for detecting any two-base insertion nearby. These data indicated that the insertion detection error rates were extremely low. ISC5 strains showed higher insertion error rate but these were likely associated with base errors in the detection of the AQP1 insertion. **E. The sequence depth and coverage statistics**: The average values of median depth of 36 chromosomes, the depth standard deviation, the genome coverage of SureSelect enriched and raw samples were given for each sample. Low sequence depth samples were marked as LOWDEP, indicating that these samples were not included in the main phylogenetic tree because of their missing depth and coverage. Somy detection accuracy classification was assigned to each sample. High corresponds to 36 high genome coverage samples, Medium corresponds to 16 medium range genome coverage samples and Low corresponds to 9 low genome coverage samples. NA corresponds to 2 samples with almost no informative genome coverage. Further descriptions were found in the text. **F. Sympatric genotypic diversity at VDC (village development committee, the lowest administrative unit in Nepal) level (available on request to authors).** VDCs where 2 different ISC genotypes (or more) were identified. See more details on the corresponding samples in S1 Table 1. *, because of lower sequence quality, ISC genotype was defined by a set of diagnostic SNPs. ^1^ Patients 80, 207 and 208 belong to the same family: they presented 2 different genotypes sampled at 10 months of interval. ^2^ Patients 89 and 91 belong to the same family: they presented 2 different genotypes sampled at the same time. **G. New insertions identified in SureSelect samples**: These indels were in non-repetitive regions and supported with depth greater than 60. They were only identified in SureSelect samples. This illustrated that previously unknown indels can be detected in SureSelect samples when there was sufficient depth. These indel calls were verified on the IGV alignment viewer to ensure the accuracy of indel alternative allele frequencies. *(S1Dataset.xlsx)*

## References

1. Burza S., Croft S.L., Boelaert M. (2018). Leishmaniasis. Lancet 392:951-970. https://doi.org/10.1016/S0140-6736(18)31204-2

2. Alvar J., Vélez I.D., Bern C., Herrero M., Desjeux P., Cano J. et al. (2012). Leishmaniasis worldwide and global estimates of its incidence. PLoS One 7:e35671. https://doi.org/10.1371/journal.pone.0035671

3. Rijal S., Sundar S., Mondal D., Das P., Alvar J., Boelaert M. (2019). Eliminating visceral leishmaniasis in South Asia: the road ahead. British Medical Journal 364:k5224. https://www.bmj.com/content/364/bmj.k5224

4. Dujardin J.C., Decuypere S. (2013). Epidemiology of leishmaniasis in the time of drug resistance. In: Drug Resistance in Leishmania Parasites. Consequences, Molecular Mechanism and Possible Treatments. Eds Ponte-Sucre A, Padron-Nieves M, Diaz E. Spinger-Verlag, Wienen. pp 65-83. https://doi.org/10.1007/978-3-7091-1125-3_4

5. Dujardin J.C. (2018). Epidemiology of leishmaniasis in the time of drug resistance (the Miltefosine era). In: Drug resistance in Leishmania parasites: consequences, molecular mechanisms and possible treatments. Eds Ponte-Sucre A & Padron-Nieves M. Springer International Publishing, pp 85–107. https://doi.org/10.1007/978-3-319-74186-4_4

6. Cuypers B., Berg M., Imamura H., Dumetz F., De Muylder G., Domagalska M.A. et al. (2018). Integrated genomic and metabolomic profiling of ISC1, an emerging *Leishmania donovani* population in the Indian subcontinent. Infection Genetics and Evolution 62:170–178. https://doi.org/10.1016/j.meegid.2018.04.021

7. Karunaweera N.D., Ferreira M.U. (2018). Leishmaniasis: current challenges and prospects for elimination with special focus on the South Asian region. Parasitology. 145:425–429. https://doi.org/10.1017/S0031182018000471

8. Imamura H., Downing T., Van den Broeck F., Sanders M.J., Rijal S., Sundar S., et al. (2016). Evolutionary genomics of epidemic visceral leishmaniasis in the Indian subcontinent. Elife 5:e12613. https://doi.org/10.7554/eLife.12613.001

9. Dumetz F., Cuypers B., Imamura H., Zander D., D’Haenens E., Maes I. et al. (2018). Molecular Preadaptation to Antimony Resistance in *Leishmania donovani* on the Indian Subcontinent. mSphere 3:e00548–17. https://doi.org/10.1128/mSphere.00548-17

10. Ostyn B., Gidwani K., Khanal B., Picado A., Chappuis F., Singh S.P. et al. (2011). Incidence of symptomatic and asymptomatic *Leishmania donovani* infections in high-endemic foci in India and Nepal: a prospective study. PLoS Neglected Tropical Diseases 5:e1284. https://doi.org/10.1371/journal.pntd.0001284

11. Dumetz F., Imamura H., Sanders M., Seblova V., Myskova J., Pescher P. et al. (2017). Modulation of Aneuploidy in *Leishmania donovani* during Adaptation to Different In Vitro and In Vivo Environments and Its Impact on Gene Expression. MBio 8:e00599–17. https://doi.org/10.1128/mBio.00599-17

12. Cuypers B. (2018). A systems biology approach for a comprehensive understanding of molecular adaptation in *Leishmania donovani*. PhD thesis, University of Antwerp. https://anet.be/record/opacuantwerpen/c:lvd:14740884/E

13. Feehery G.R., Yigit E., Oyola S.O., Langhorst B.W., Schmidt V.T., Stewart F.J. et al. (2013). A method for selectively enriching microbial DNA from contaminating vertebrate host DNA. PLoS One 8:e76096. https://doi.org/10.1371/journal.pone.0076096

14. Oyola S.O., Ariani C.V., Hamilton W.L., Kekre M., Amenga-Etego L.N., Ghansah A., et al. (2016). Whole genome sequencing of *Plasmodium falciparum* from dried blood spots using selective whole genome amplification. Malaria Journal 15:597. https://doi.org/10.1186/s12936-016-1641-7

15. Doyle R.M., Burgess C., Williams R., Gorton R., Booth H., Brown J. et al. (2018). Direct Whole-Genome Sequencing of Sputum Accurately Identifies Drug-Resistant *Mycobacterium tuberculosis* Faster than MGIT Culture Sequencing. Journal of Clinical Microbiology 56:e00666-18. https://doi.org/10.1128/JCM.00666-18

16. Kozarewa I., Ning Z., Quail M.A., Sanders M.J., Berriman M., Turner D.J. (2009). Amplification-free Illumina sequencing-library preparation facilitates improved mapping and assembly of (G+C)-biased genomes. Nature Methods 6:291-5. https://doi.org/10.1038/nmeth.1311

17. Bronner I.F., Quail M.A., Turner D.J., Swerdlow H. (2014). Improved Protocols for Illumina Sequencing. Current Protocols of Human Genetics 80:18.2.1-42. https://doi.org/10.1002/0471142905.hg1802s80

18. Downing T., Imamura H., Decuypere S., Clark T.G., Coombs G.H., Cotton J.A. et al. (2011). Whole genome sequencing of multiple *Leishmania donovani* clinical isolates provides insights into population structure and mechanisms of drug resistance. Genome Research 21:2143-56. https://doi.org/10.1101/gr.123430.111

19. Tamura K., Peterson D., Peterson N., Stecher G., Nei M., Kumar S. (2011). MEGA5: molecular evolutionary genetics analysis using maximum likelihood, evolutionary distance, and maximum parsimony methods. Molecular Biology and Evolution 28:2731-9. https://doi.org/10.1093/molbev/msr121

20. Huson D.H., Bryant D. (2006). Application of phylogenetic networks in evolutionary studies. Molecular Biology and Evolution 23:254-67. https://doi.org/10.1093/cid/cit102

21. Tobie, E. J., Von Brand, T. & Mehlman, B. (1950). Cultural and physiological observations on *Trypanosoma rhodesiense* and *Trypanosoma gambiense*. Journal of Parasitology 36:48-54.

22. Yardley V., Croft S.L., De Doncker S., Dujardin J.C., Koirala S., Rijal S. et al. (2005). The sensitivity of clinical isolates of *Leishmania* from Peru and Nepal to miltefosine. American Journal of Tropical Medicine and Hygiene 73:272-5.

23. Rai K., Bhattarai N.R., Vanaerschot M., Imamura H., Gebru G., Khanal B. et al. (2017). Single locus genotyping to track *Leishmania donovani* in the Indian subcontinent: Application in Nepal. PLoS Neglected Tropical Diseases 11:e0005420. https://doi.org/10.1371/journal.pntd.0005420

24. Rogers M.B., Hilley J.D., Dickens N.J., Wilkes J., Bates P.A., Depledge D.P. et al. (2011). Chromosome and gene copy number variation allow major structural change between species and strains of *Leishmania*. Genome Research 21:2129-42. https://doi.org/10.1101/gr.122945.111

25. Prieto Barja P., Pescher P., Bussotti G., Dumetz F., Imamura H., Kedra D. et al. (2017). Haplotype selection as an adaptive mechanism in the protozoan pathogen *Leishmania donovani*. Nature Ecology and Evolution 1:1961-1969. https://doi.org/10.1038/s41559-017-0361-x

26. Pacheco M.A., Lopez-Perez M., Vallejo A.F., Herrera S., Arévalo-Herrera M., Escalante A.A. (2016). Multiplicity of Infection and Disease Severity in *Plasmodium vivax*. PLoS Neglected Tropical Diseases 10:e0004355. https://doi.org/10.1371/journal.pntd.0004355

27. Llewellyn M.S., Messenger L.A., Luquetti A.O., Garcia L., Torrico F., Tavares S.B. et al. (2015). Deep sequencing of the *Trypanosoma cruzi* GP63 surface proteases reveals diversity and diversifying selection among chronic and congenital Chagas disease patients. PLoS Neglected Tropical Diseases 9:e0003458. https://doi.org/10.1371/journal.pntd.0003458

28. Sánchez L.V., Bautista D.C., Corredor A.F., Herrera V.M., Martinez L.X., Villar J.C. et al. (2013). Temporal variation of *Trypanosoma cruzi* discrete typing units in asymptomatic Chagas disease patients. Microbes and Infection 15:745-8. https://doi.org/10.1016/j.micinf.2013.06.008

29. Ferreira Ede C., Cruz I., Cañavate C., de Melo L.A., Pereira A.A., Madeira F.A., Valério S.A. et al. (2015). Mixed infection of *Leishmania infantum* and *Leishmania braziliensis* in rodents from endemic urban area of the New World. BMC Veterinary Research 11:71. https://doi.org/10.1186/s12917-015-0392-y

30. Soares I.R., Silva S.O., Moreira F.M., Prado L.G., Fantini P., Maranhão Rde P. et al. (2013). First evidence of autochthonous cases of *Leishmania (Leishmania) infantum* in horse (Equus caballus) in the Americas and mixed infection of *Leishmania infantum* and *Leishmania (Viannia) braziliensis*. Veterinary Parasitology 197:665-9. https://doi.org/10.1016/j.vetpar.2013.06.014

31. Veland N., Valencia B.M., Alba M., Adaui V., Llanos-Cuentas A., Arevalo J., et al. (2013). Simultaneous infection with *Leishmania (Viannia) braziliensis* and *L. (V.) lainsoni* in a Peruvian patient with cutaneous leishmaniasis. American Journal of Tropical Medicine and Hygiene 88:774-7 https://doi.org/10.4269/ajtmh.12-0594

32. Shirian S., Oryan A., Hatam G.R., Daneshbod Y. (2012). Mixed mucosal leishmaniasis infection caused by *Leishmania tropica* and *Leishmania major*. Journal of Clinical Microbiology 50:3805-8. https://doi.org/10.1128/JCM.01469-12

33. Zackay A., Cotton J.A., Sanders M., Hailu A., Nasereddin A., Warburg A. et al. (2018). Genome wide comparison of Ethiopian *Leishmania donovani* strains reveals differences potentially related to parasite survival. PLoS Genetics 14:e1007133. https://doi.org/10.1371/journal.pgen.1007133

34. Pratlong F., Portus M., Rispail P., Moreno G., Bastien P., Rioux J.A. (1989). Simultaneous presence in dogs of 2 zymodemes of the *Leishmania infantum* complex. Annales de Parasitologie Humaine et Comparée 64:312-4. https://doi.org/10.1051/parasite/1989644312

35. Rogers M.B., Downing T., Smith B.A., Imamura H., Sanders M., Svobodova M. et al. (2014). Genomic confirmation of hybridisation and recent inbreeding in a vector-isolated *Leishmania* population. PLoS Genetics 10:e1004092. https://doi.org/10.1371/journal.pgen.1004092

36. Odiwuor S., De Doncker S., Maes I., Dujardin J.C., Van der Auwera G. (2011). Natural *Leishmania donovani/Leishmania aethiopica* hybrids identified from Ethiopia. Infection Genetics and Evolution 11:2113-8. https://doi.org/10.1016/j.meegid.2011.04.026

