## Supplemental Text for "Genomes of *Leishmania* parasites directly sequenced from patients with visceral leishmaniasis in the Indian subcontinent"

**S1 Text**

1. Supplementary results

**SureSelect bait design**

To sequence the genome of *Leishmania donovani* directly in clinical samples collected from VL patients, we developed a method to enrich *Leishmania* nuclear DNA from samples containing mostly human DNA. The method is based on a custom Agilent SureSelect array with baits designed to tile the *L. donovani* BPK282 assembly based on 454 and Illumina sequencing data [18], including contigs not assembled into chromosomes. Baits were tiled 'end to end' with no gaps or overlaps against this reference. Additional probes were added to capture regions of *L. infantum* JPCM5 [11] Leishmania_infantum_JPCM5_27July2011.fa adding probes against regions not covered by BPK282 probes with less than 5bp mismatching. Probes were subsequently removed from this design if they overlapped a mask of repetitive or low complexity sequence in BPK282 by at least 40bp, or if they matched reference assemblies of mouse, human or dogs with at least 40bp identity. The final design contained 218,904 RNA probes of 120bp, covering 26 Mbp of the 32 Mb genome. The inclusion of additional probes designed to capture both species, and subsequent changes to the genome assembly, resulted in some small regions of the genome being covered by more than one probe (see S1 Fig. A).

**SureSelect genome capture and sequencing (SuSL-seq) optimization on artificial mixtures**

We tested the baits on a series of artificial mixtures of DNA isolated from axenic cultured *Leishmania* promastigotes and commercially obtained human and mouse genomic DNA. Artificial mixtures were prepared using commercially available mouse or human DNA

(Promega) and parasite DNA extracted from cultured promastigotes (BPK282 or BPK026). These mixtures ("artificial mixtures") were produced with 3 different percentages: 0.06%, 0.006% and 0.0006%, which encompassed the range of parasite loads reported in bone marrow samples collected in VL patients in the ISC (Ref S1). Overall, the *Leishmania* genome was captured and sequenced evenly with exceptions in some repetitive regions (mini-exon, the ends of chromosome 1) that were over-represented in SureSelect libraries. The highest coverage was found in the genomic regions targeted with baits (S1 Fig. B), but sequence reads were also observed over many regions without probes, as DNA fragments included in the sequencing library fragments extended beyond the probes by up to several hundred basepairs (S1 Fig. C). After application of the baits, we obtained enrichment in the range of 400–1,100 fold, with the highest enrichment in the samples with lowest Leishmania DNA percentage (likely due to saturation of baits for higher percentages) (S2 Fig. A-B).

We further investigated the relationship between read depth, genome coverage and the initial Leishmania DNA percentage in a sample. After normalizing the sequence outputs, followed by taking random sub-samples from the pooled reads across all 4 replicates for each Leishmania DNA percentage, we calculated how the percentage of bases covered at read depths (1, 5, 10, 20, 30, 50, and 100) varied with the total number of reads (S2 Fig. C). For the lowest *Leishmania* DNA percentage (0.0006%), increasing the total read number did not improve genome coverage, with only ca. 30% of genome being covered at least 1x, and almost no sites covered to higher depths. We concluded from this that samples with Leishmania DNA percentages below 0.006% are not suitable for whole genome-based analysis with the current method. For higher Leishmania DNA percentages, it was clear that increasing the total number of reads improved the quality of the results: with 104 million reads, approximately 20% and

90% genome was covered at least 5x, for the 0.006% and 0.06% samples, respectively (S2 Fig. C).

We next evaluated whether these samples were suitable for analysis of genetic diversity, by testing which types of genetic variants could be reliably called from SureSelect libraries. We could correctly measure aneuploidy in the samples with highest Leishmania DNA percentages (0.06%), while this was impossible for the 0.0006% sample. At 0.006%, we could not precisely assess somy of the chromosomes, but we were able to detect that chromosome 31, and longer chromosomes (Chromosomes 23-35) had a somy higher than 2 (S2 Fig. D). Single nucleotide polymorphism (SNP) calling on the artificial mixtures with high Leishmania DNA percentages identified essentially identical variants in these samples in comparison to pure DNA of the corresponding isolate, differing by 6 to 9 heterozygous bases out of 197 variant sites.

**Direct sequencing of *Leishmania donovani* genomes in 63 clinical samples**

In our protocol, an indexed multiplex library is generated prior to SureSelect capture. This makes the SureSelect procedure more cost-effective, as multiple samples can be processed in a single SureSelect experiment, but to ensure sufficient representation of every sample in the final (post-SureSelect) library pool, samples need to have similar levels of target (i.e. *Leishmania*) DNA before pooling. To ensure this, and to precisely quantify the input libraries, the amount of *Leishmania* DNA present in the samples was measured by quantitative PCR (qPCR). The qPCR method applied here was validated in the first set of 24 samples by deep sequencing without SureSelect enrichment and comparing the estimated amounts of Leishmania DNA obtained by both methods (S4 Fig. B). We selected samples with *Leishmania*

DNA percentages of at least 0.006%, and the samples were processed in total of 6 batches in 2 rounds, where 500ng and 100ng of total DNA were respectively used to prepare sequencing libraries. The sequencing statistics for all clinical samples are summarized in S1 Dataset E.

### 2. Supplementary methods

#### Clinical samples

For 12 clinical samples, we had paired isolates from the same patients: bone marrow aspirate was diluted in 3.5 ml Locke solution and split in two. First, 0.5 ml was put in a PCR-tube that was directly frozen at -70°C and used in this study for SuSL analysis. Secondly, 3 ml were put in Tobie culture medium for isolation attempt and cryopreservation as soon as the culture was positive. These promastigotes had been propagated in vitro for 12-22 passages. Details are presented in S1 Table and geographical origin of samples is visualized in S3 Fig. Laboratory methods, clinical definitions and procedures are described elsewhere [8,18, Ref S2-Ref S3].

All samples here analyzed were collected on admission, thus before the onset of therapy at BPKIHS (in a few exceptions, some patients had already received a treatment previously, see remarks in S1 Table). A rigorous attention was paid to the coding of samples, according to the procedures implemented at BPKIHS. Briefly, each patient received a unique VL (VL-project) or BPK (Leishnatdrug-R, Kaladrug-R and SINGLE) number on admission. When a culture was successfully derived from a given patient, the same BPK number was used: thus for instance, from patient BPK157, the isolate MHOM/NP/02/BPK157/0 (/0 standing for an isolation made before the onset of treatment) was derived. In current paper, we used the suffix BM or SP to identify bone marrow or spleen samples, respectively.

The % of Leishmania DNA in clinical samples was estimated using real-time quantitative PCR (qPCR). The qPCR was carried out a LightCycler480 (Roche, Basel, Switzerland) using SensiMix SYBR No-ROX (Bioline, London, UK) in 25µl reaction volume using primers specific to a single copy *Leishmania* locus, Cysteine synthase (CS, Ref S4) and to a single locus in human genome, RPL30 (NEB Next Microbiome Enrichment kit, NEB, Ipswich, USA). The following cycling conditions were used: 95˚C for 10 mins, followed by 45 rounds of 95˚C for 30s, 60˚C for 15s, 72˚C for 15s. On each reaction plate, standard curves for *Leishmania*, prepared from genomic DNA of BPK282 or BPK026 strain and for human DNA prepared from commercially available human genomic DNA (Promega, Fitchburg, USA) were included and were used to calculate the amounts of respective DNA in clinical samples.

**Amplicon sequencing**

The following primers were used to amplify loci containing a diagnostic SNP/insertion specific of ISC groups 3, 4, 5 and 6:

ISC3_for_NGS ACACTCTTTCCCTACACGACGCTCTTCCGATCTTGCGGGGTGAAAGGCATCCTG,

ISC3_rev_NGS TCGGCATTCCTGCTGAACCGCTCTTCCGATCTCATGCCGGACACTGACCCGT,

ISC4_for_NGS ACACTCTTTCCCTACACGACGCTCTTCCGATCTTCCAACCCTCGTGGCGCAAG,

ISC4_rev_NGS TCGGCATTCCTGCTGAACCGCTCTTCCGATCTACACCCACACCGTGCGATCC,

ISC5_for_NGS ACACTCTTTCCCTACACGACGCTCTTCCGATCTAGGTGTTTGCAACATTAGGGTA,

ISC5_rev_NGS TCGGCATTCCTGCTGAACCGCTCTTCCGATCTTGTACAAATACCGATGGTGGCT,

ISC6_for_NGS ACACTCTTTCCCTACACGACGCTCTTCCGATCTGCTCCCCTTCACTGCACTCCC,

ISC6_rev_NGS TCGGCATTCCTGCTGAACCGCTCTTCCGATCTAGGGTGGAGAGAATGTCAACGCA.

For each ISC group specific PCR, we used DNA amplified from strains, which served as positive, negative (lack of diagnostic SNP/insertion) controls. For ISC3, BPK067 was used as a positive control, and BPK282, and BPK275 as negative controls. For ISC4, BPK206, BPK080 were used as a positive controls, and BPK282, and BPK275 as negative controls. For ISC5, BPK275, BPK173 were used as a positive controls, and BPK282, and BPK206 as negative controls. For ISC6, BPK282, BPK085 were used as a positive control, and BPK275, and BPK206 as negative controls.

### 3. Supplementary references

Ref S1. Verma S., Kumar R., Katara G.K., Singh L.C., Negi N.S., Ramesh V. et al. (2010). Quantification of parasite load in clinical samples of leishmaniasis patients: IL-10 level correlates with parasite load in visceral leishmaniasis. *PLoS One* 5:e10107. https://doi.org/10.1371/journal.pone.0010107

Ref S2. De Doncker S., Hutse V., Abdellati S., Rijal S., Singh Karki B.M., Decuypere S. et al. (2005). A new PCR-ELISA for diagnosis of visceral leishmaniasis in blood of HIV-negative subjects. *Transactions of the Royal Society of Tropical Medicine and Hygiene* 99:25-31. https://doi.org/10.1016/j.trstmh.2004.01.015

Ref S3. Rijal S., Ostyn B., Uranw S., Rai K., Bhattarai N.R., Dorlo T.P. et al. (2013). Increasing failure of miltefosine in the treatment of Kala-azar in Nepal and the potential role of parasite drug resistance, reinfection, or noncompliance. *Clinical Infectious Diseases* 56:1530-8. https://doi.org/10.1093/cid/cit102

Ref S4. Decuypere S., Vanaerschot M., Rijal S., Yardley V., Maes L., De Doncker S. et al. (2008). Gene expression profiling of *Leishmania (Leishmania) donovani*: overcoming technical variation and exploiting biological variation. *Parasitology* 135:183-194. https://doi.org/10.1017/S0031182007003782
