## Supplementary figures for "Genomes of *Leishmania* parasites directly sequenced from patients with visceral leishmaniasis in the Indian subcontinent"

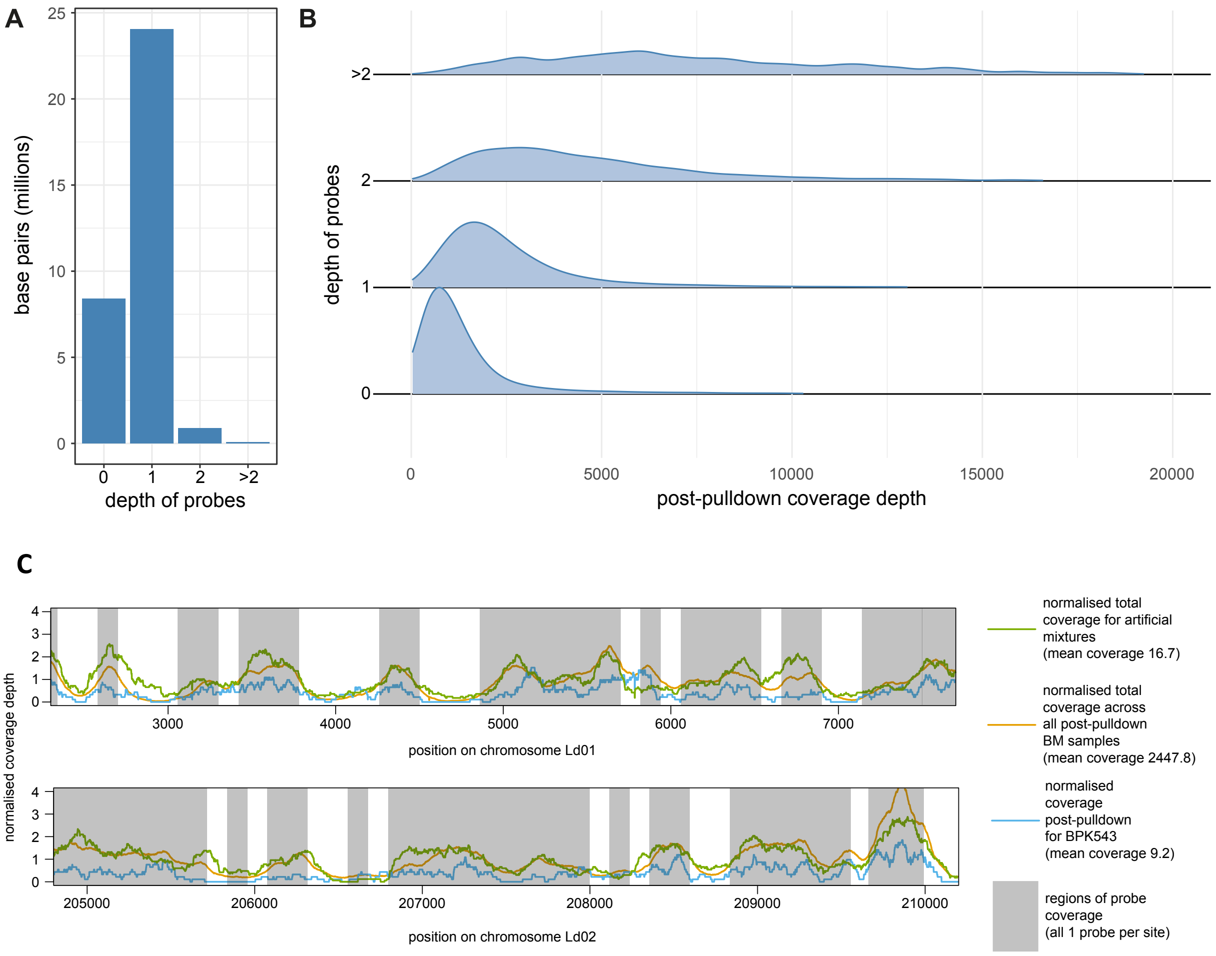


**Fig. S1. A.** Distribution of probes across *L. donovani* BPK282 reference genome. Most of the reference genome is covered by only a single probe. Some regions are not covered to avoid repetitive regions and regions with homology to mammalian hosts. Changes in the genome assembly subsequent to the bait design and the presence of additional probes to capture *L. infantum* have led to some regions being covered by multiple probes sequences. **B.** Much of the variation in depth in coverage in clinical samples is due to the presence of multiple probe sequences. Plots show the total coverage density across all clinical samples for regions of the reference genome covered by 0, 1, 2 or more probe sequences. **C.** Depth of read coverage in SureSelect enriched samples across two regions of the *Leishmania* genome. Lines show normalized coverage depth (actual coverage per base pair divided by genome-wide mean coverage) for one clinical sample (BPK543, blue) and the sum for all clinical samples (orange). Grey shading shows regions where SureSelect probes map to the genome. Mean coverage for BPK543 was 9.2 reads, for the summed clinical samples 2447.8.

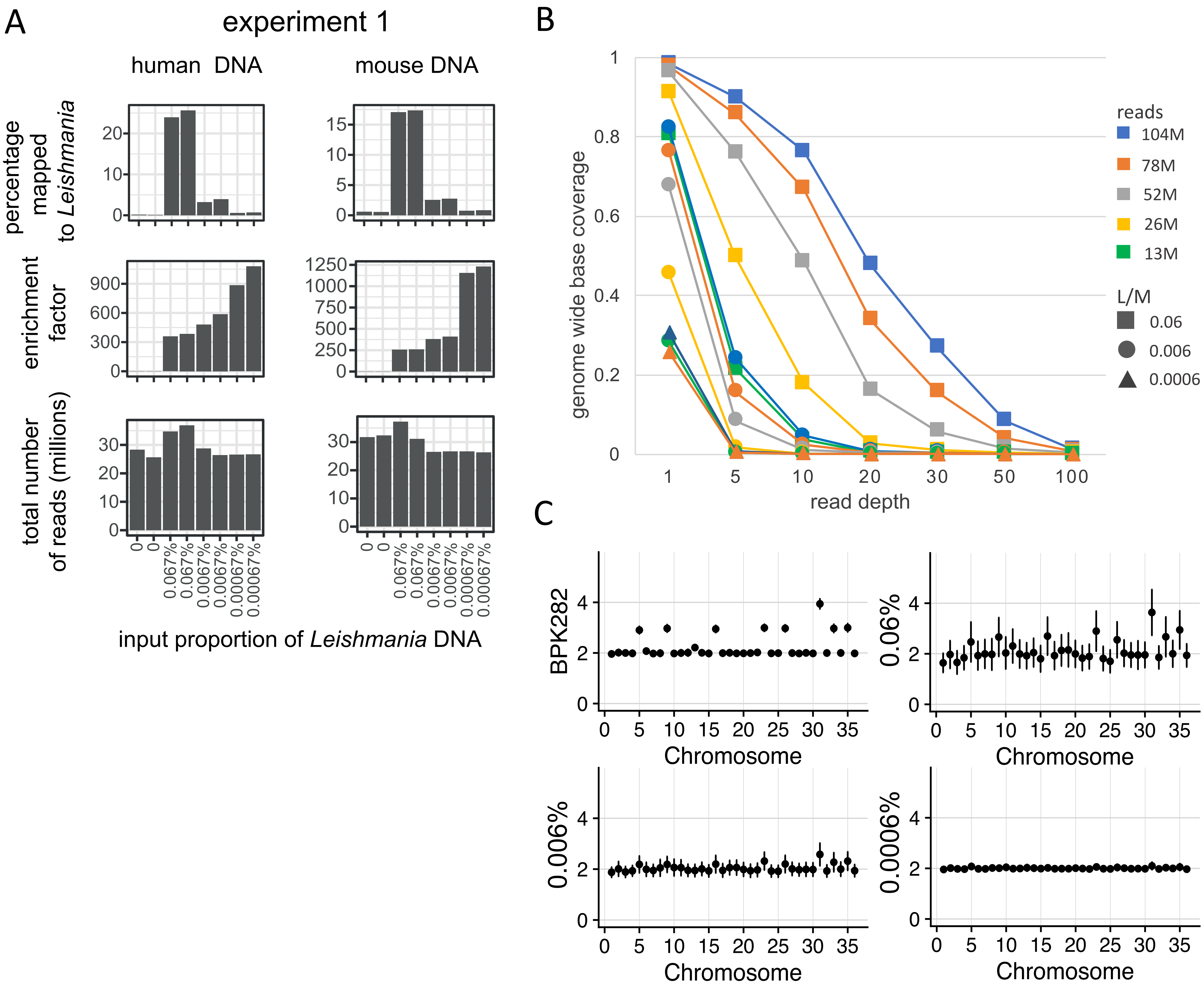


**Fig. S2. Performance of SuSL-sequencing on artificial mixtures of *Leishmania* and mammal DNA at three different *Leishmania* DNA percentages: 0.06, 0.006 and 0.0006 %. A**. Summary statistics for sequencing data from these experiments, showing the total number of reads in each library, the proportion of reads mapping to the *L. donovani* reference genome, and the enrichment factor, calculated as the ratio of the proportion of reads mapping to *L. donovani* in the SureSelect libraries to the proportion of *Leishmania* promastigote DNA included in the pre-pulldown DNA mixtures. **B.** Evenness of genome coverage. The y-axis shows the proportion of bases covered with a minimum number of sequencing reads -- read depth along the x-axis -- by reads extracted from SureSelect sequencing data derived from artificial mixtures. 104M, 78M, 52M, 26M and 13M indicate the total number of reads in millions sampled from each library; 0.06, 0.006 and 0.0006 are the input proportions of *Leishmania* DNA in percent. For lower *Leishmania* DNA percentages, the higher read depth did not result in higher genome base coverage. **C**. Inferred somy from SureSelect on artificial mixtures. BPK282, pure DNA from promastigote; LD06-0006, simulated clinical samples at three *Leishmania* DNA percentages 0.06%, 0.006% and 0.0006%, respectively. The y-axis shows normalized somy estimate for each chromosome along the x-axis. Points show central estimate and bars show one standard deviation around these estimates from four replicates of each *Leishmania* DNA percentages.

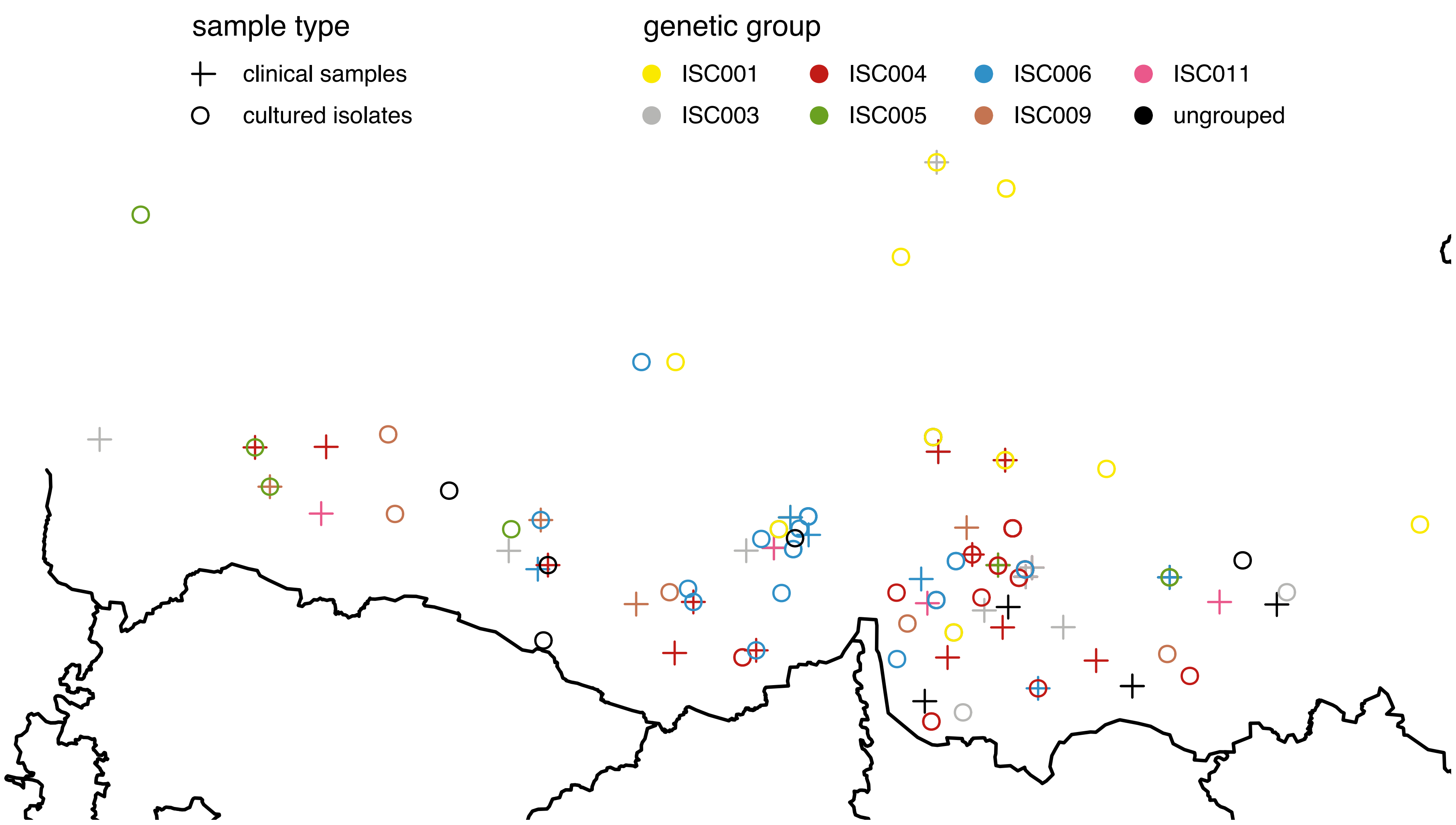


**Fig. S3. Map of Nepal showing geographical origin of clinical samples and cultured promastigote isolates.** Colours represent different genotype groups defined elsewhere (1). Circles represent Nepalese promastigote isolates from ref 1, crosses bone marrow and spleen samples.

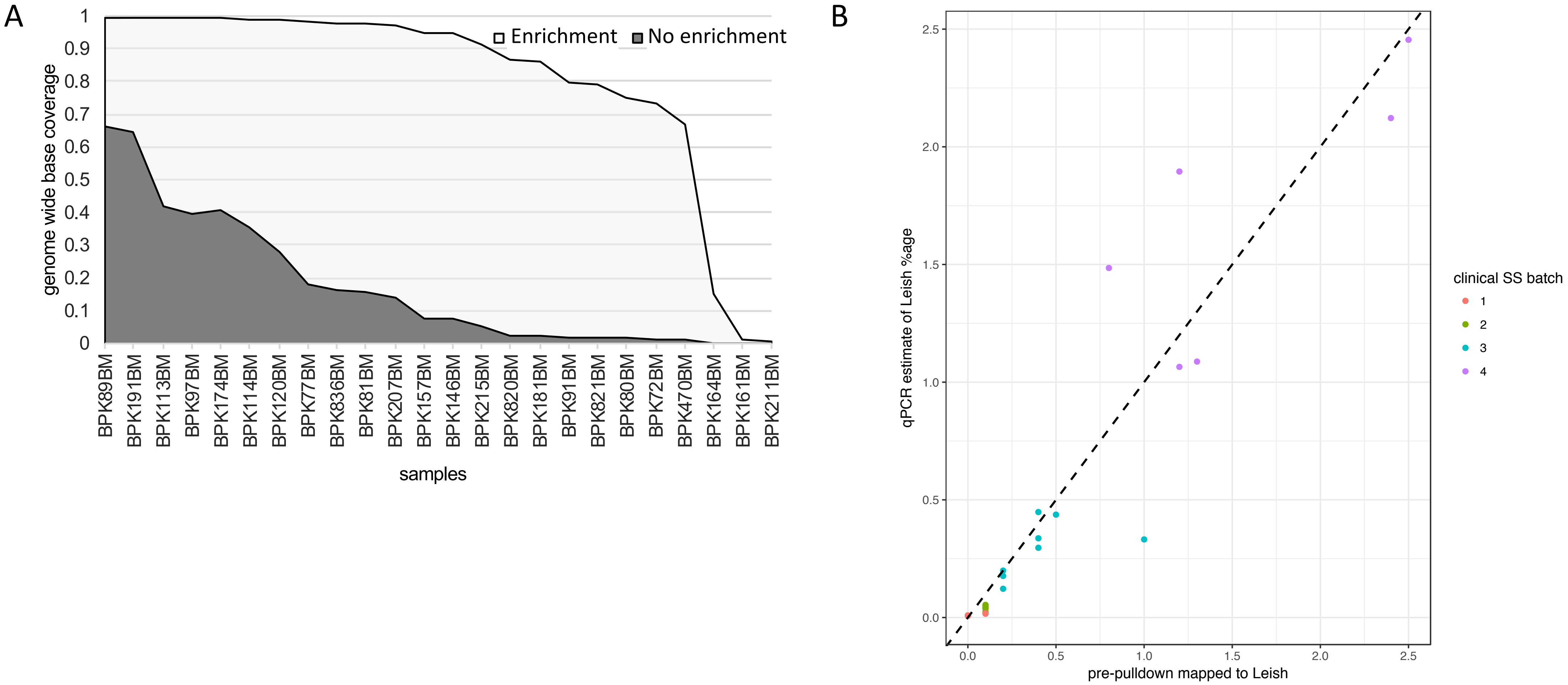


**Fig. S4. A.** Genome sequencing of clinical samples: percentage of the genome covered by more than 1 read, with and without SuSL enrichment. **B**. Relationship between qPCR estimate of *Leishmania* DNA concentration (y axis) and the proportion of sequencing reads mapping to the *L. donovani* reference genome before SureSelect enrichment in clinical samples for which both qPCR and pre-enrichment sequence data are available, confirming accuracy of the qPCR estimates. Batches were formed of samples with similar proportions of *Leishmania* DNA pre-enrichment.

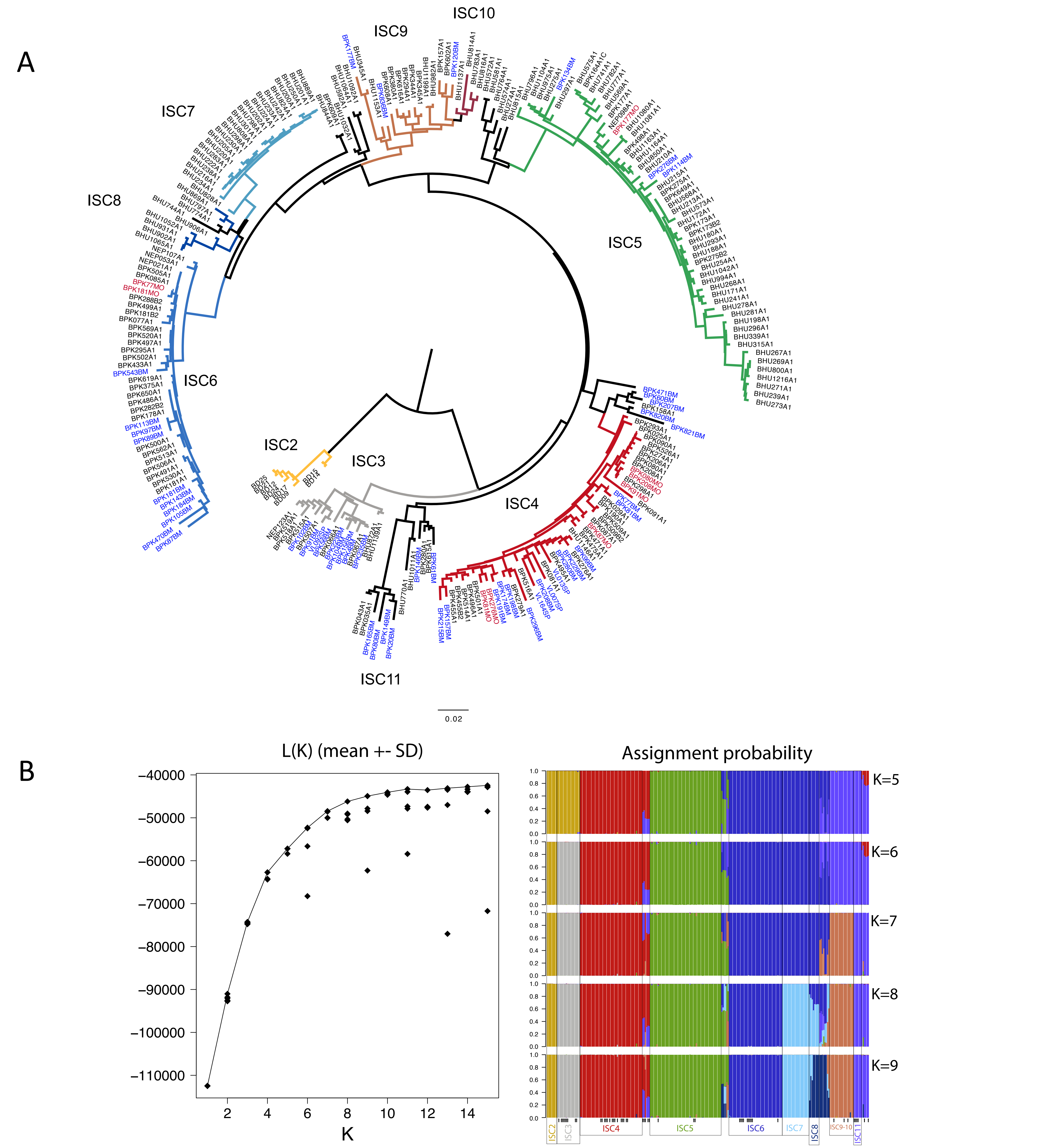


**Fig. S5. A.** Phylogenetic tree (Neighbor-joining) based on bone marrow (BM), spleen (SP), isolates paired to clinical samples (MO) and previously sequenced lines (labelled A1 or B2, ref 1). The tree is identical to the main one (Fig 2A), except that the labels of the samples are indicated. Bone marrow (BM) and spleen (SP) samples are labelled in blue; isolates that are paired to clinical samples are labelled in red, previously sequenced lines are labelled in black. ISC2-ISC10 were sub-populations previously defined (ref 1). See Fig.2A for comments on ISC11. **B.** Results from STRUCTURE analyses from five replicate runs (2x10$^6$ MCMC chains following 10$^6$ burn-in steps) under the Admixture model assuming 1-15 *K* clusters. The plot on the left shows the estimated loglikelihood of all runs for each *K* cluster. Barplots on the right summarize the assignment probabilities of every *L. donovani* sequence to each inferred cluster assuming 5, 6, 7, 8 or 9 clusters. Arrows denote the CG samples sequenced in this study.

Fig. S6. A 1

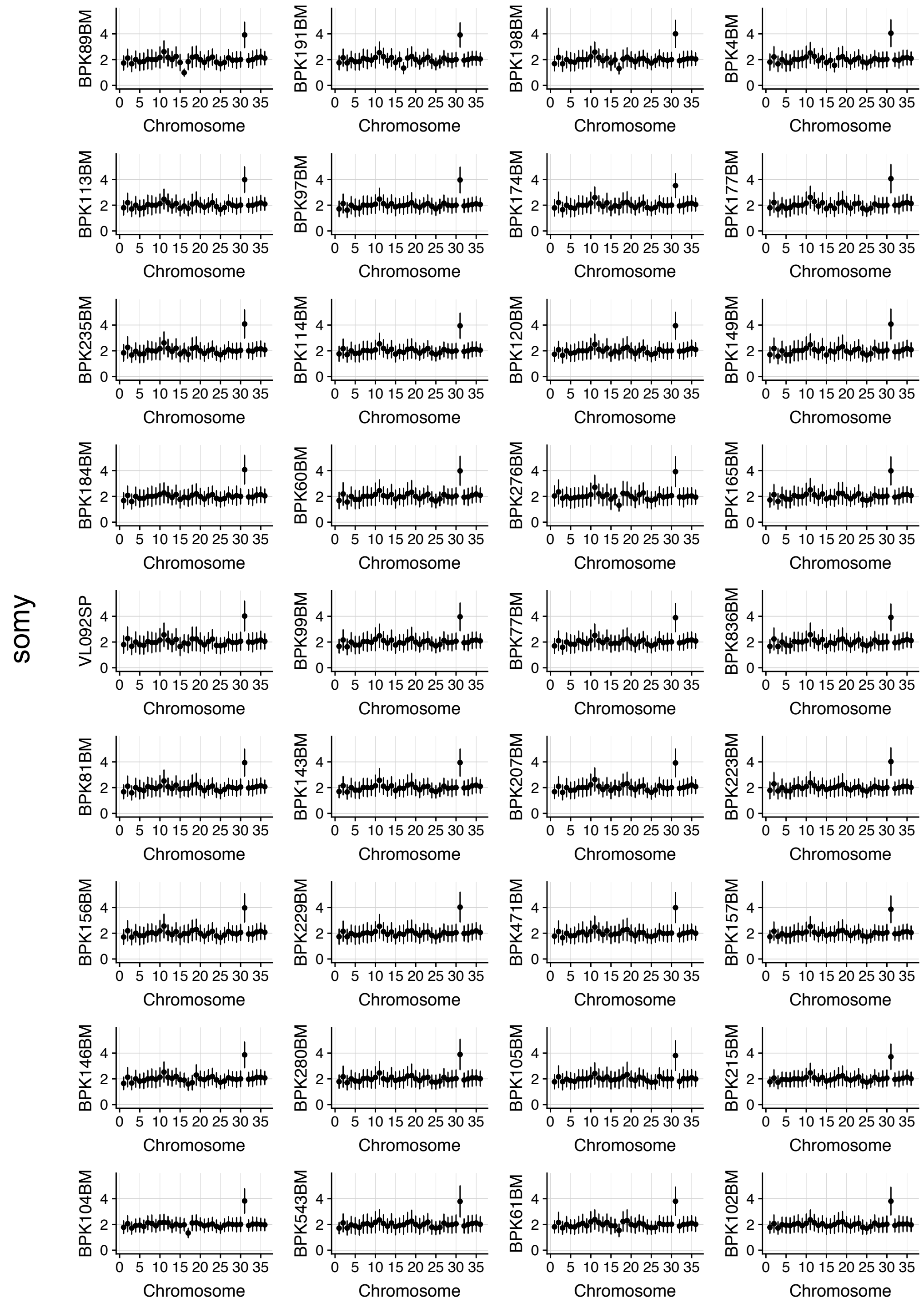

somy
BPK89BM
BPK191BM
BPK198BM
BPK4BM
BPK113BM
BPK97BM
BPK174BM
BPK177BM
BPK235BM
BPK114BM
BPK120BM
BPK149BM
BPK184BM
BPK60BM
BPK276BM
BPK165BM
VL092SP
BPK99BM
BPK77BM
BPK836BM
BPK81BM
BPK143BM
BPK207BM
BPK223BM
BPK156BM
BPK229BM
BPK471BM
BPK157BM
BPK146BM
BPK280BM
BPK105BM
BPK215BM
BPK104BM
BPK543BM
BPK61BM
BPK102BM
Chromosome
0 5 10 15 20 25 30 35
0 2 4

Fig. S6. A 2

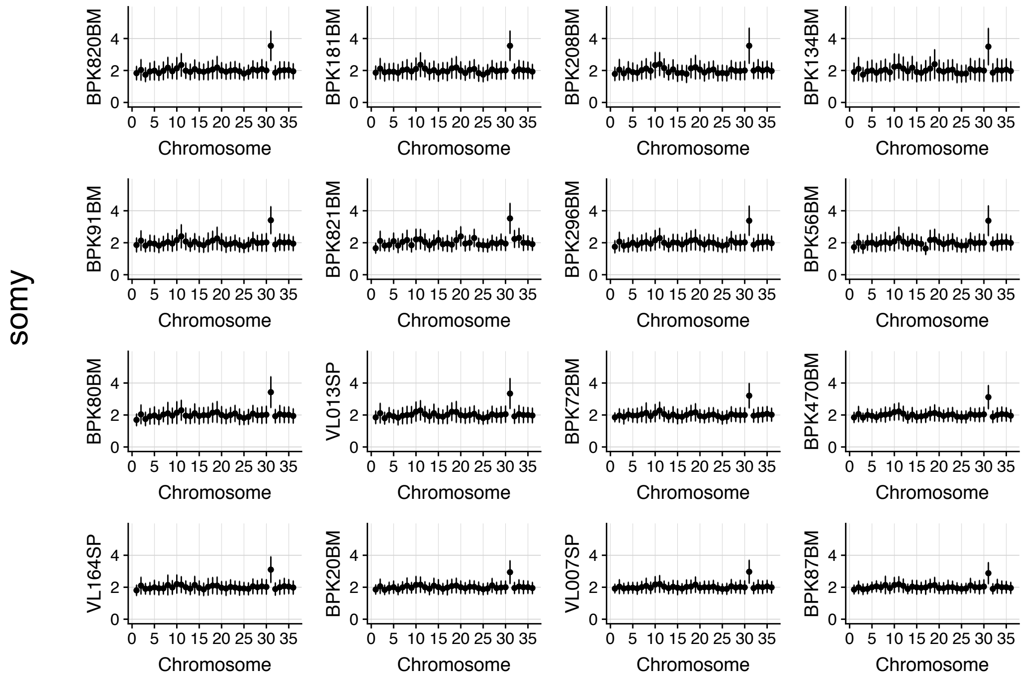


Fig. S6. A 3

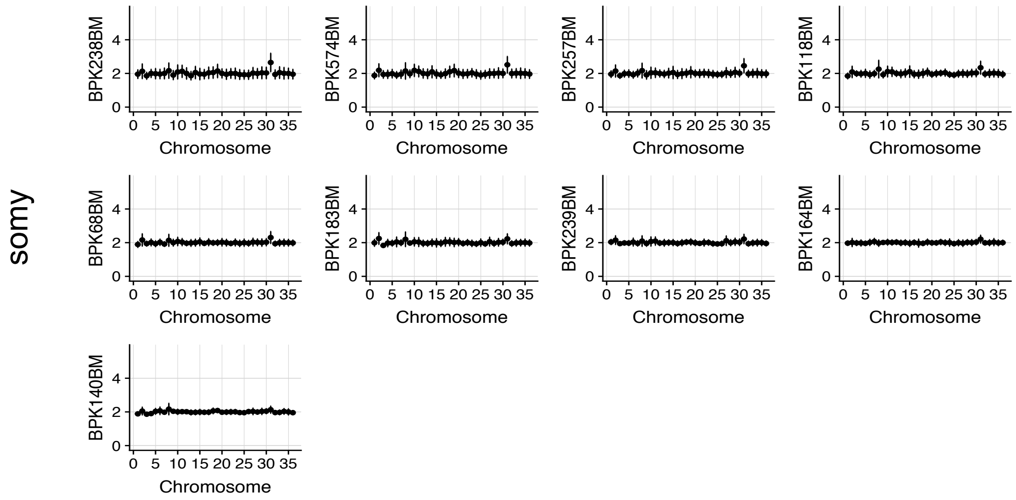

Fig. S6 B

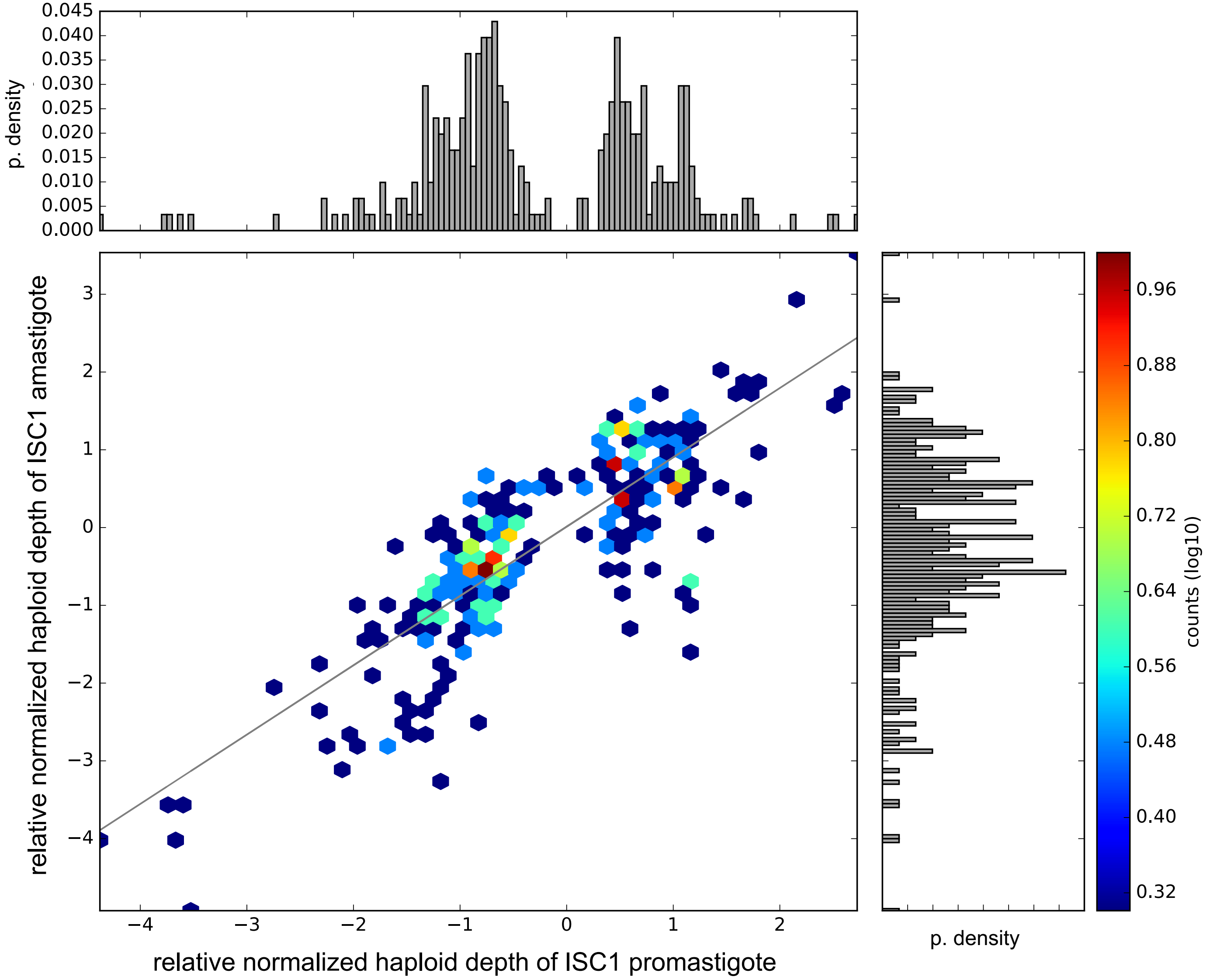

p. density
0.045
0.040
0.035
0.030
0.025
0.020
0.015
0.010
0.005
0.000
relative normalized haploid depth of ISC1 amastigote
3
2
1
0
−1
−2
−3
−4
relative normalized haploid depth of ISC1 promastigote
p. density
0.96
0.88
0.80
0.72
0.64
0.56
0.48
0.40
0.32
counts (log10)

**Fig. S6. A.** Somy in bone marrow and spleen samples. BM=Bone marrow and SP=spleen. The y-axis and x-axis represent somy and chromosome, respectively. S-values and standard deviation error bars are given in black filled circle and vertical line, respectively. The S- and standard deviation values for the bone marrow samples were based on the binned depth method. The samples were sorted in the order of higher genome coverage, the 3 categories reflect resolution of somy estimation 1) High: whose genome coverage ranging from 99.45% to 89.10%. 2) Medium: whose genome coverage ranging from 86.93% to 56.60%. 3) Low: whose genome coverage ranging from 45.80% to 9.80%. In the lower *Leishmania* DNA concentration samples, it is still possible to identify higher copy number in chromosome 31. S-values of chromosome 2 and 11 were higher for almost all samples but these higher values were also observed in artificial mixtures of BPK282 where chromosome 2 and 11 were definitively disomic. Close inspection of read depth did not reveal any particular evidences to support their S-values to be higher than 2. **B.** Correlation between gene depth at 303 ISC1-specific CNV genes identified in promastigote samples. Correlation was moderately high $r^2 = 0.68$, p-value = $10^{-76}$ and slope = 0.882 indicating that SuSL-seq can be used to confirm existing group-specific CNVs. On the plot, colored dots represent counts in log10 scale and probability density distributions were given on the corresponding axes.

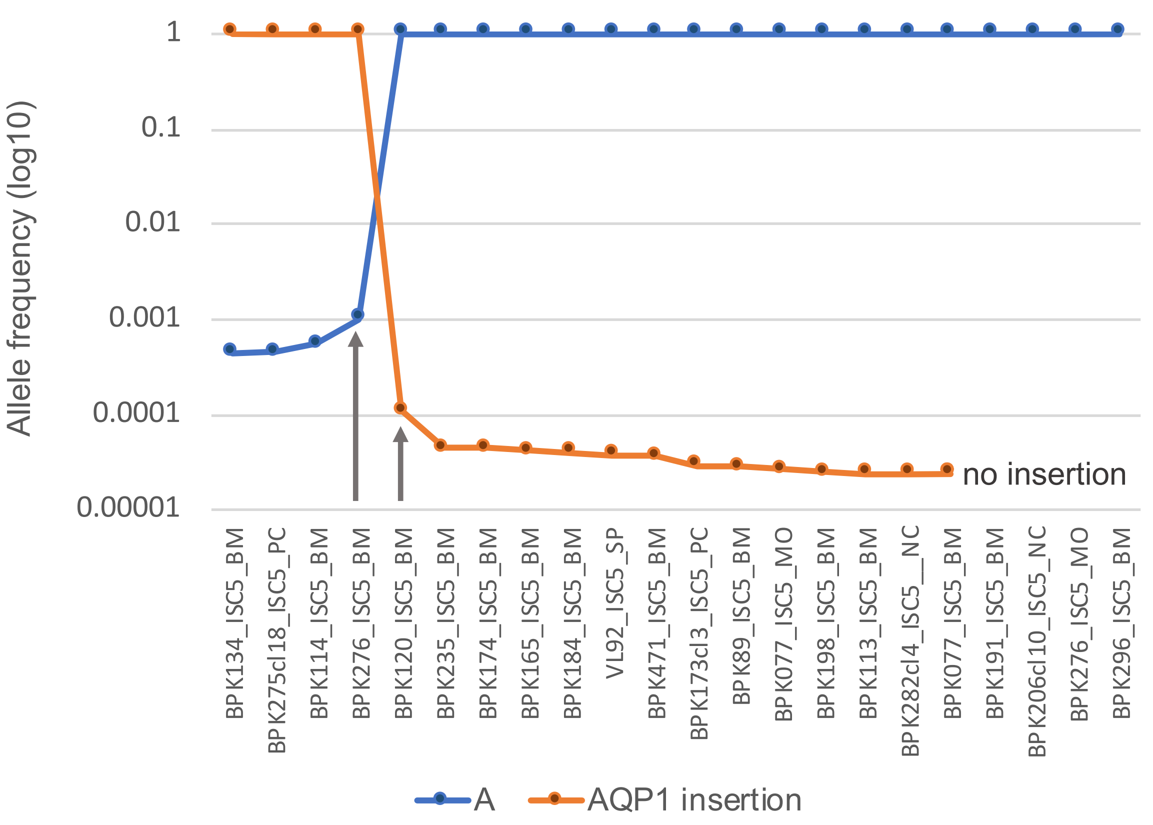


**Fig. S7. Allele frequency for the amplicon diagnostic of ISC5 genotype estimated with NGS-based deep amplicon sequencing.** Blue line represents the frequency for the non-ISC5 allele, while the orange line represents the ISC5-specific allele, which contains a 2bp-insertion in the AQP1 locus. Arrows indicate the two clinical samples in which a second allele was detected above the background, being statistically significant for BPK120_ISC5_BM. BM, Bone Marrow; SP, Spleen aspirate; MO, cultured isolate; NC, Negative Control, a cloned strain where the diagnostic ISC5-specific allele is absent; PC, Positive Control, a cloned strain characterized by the presence of the diagnostic ISC5-specific allele.

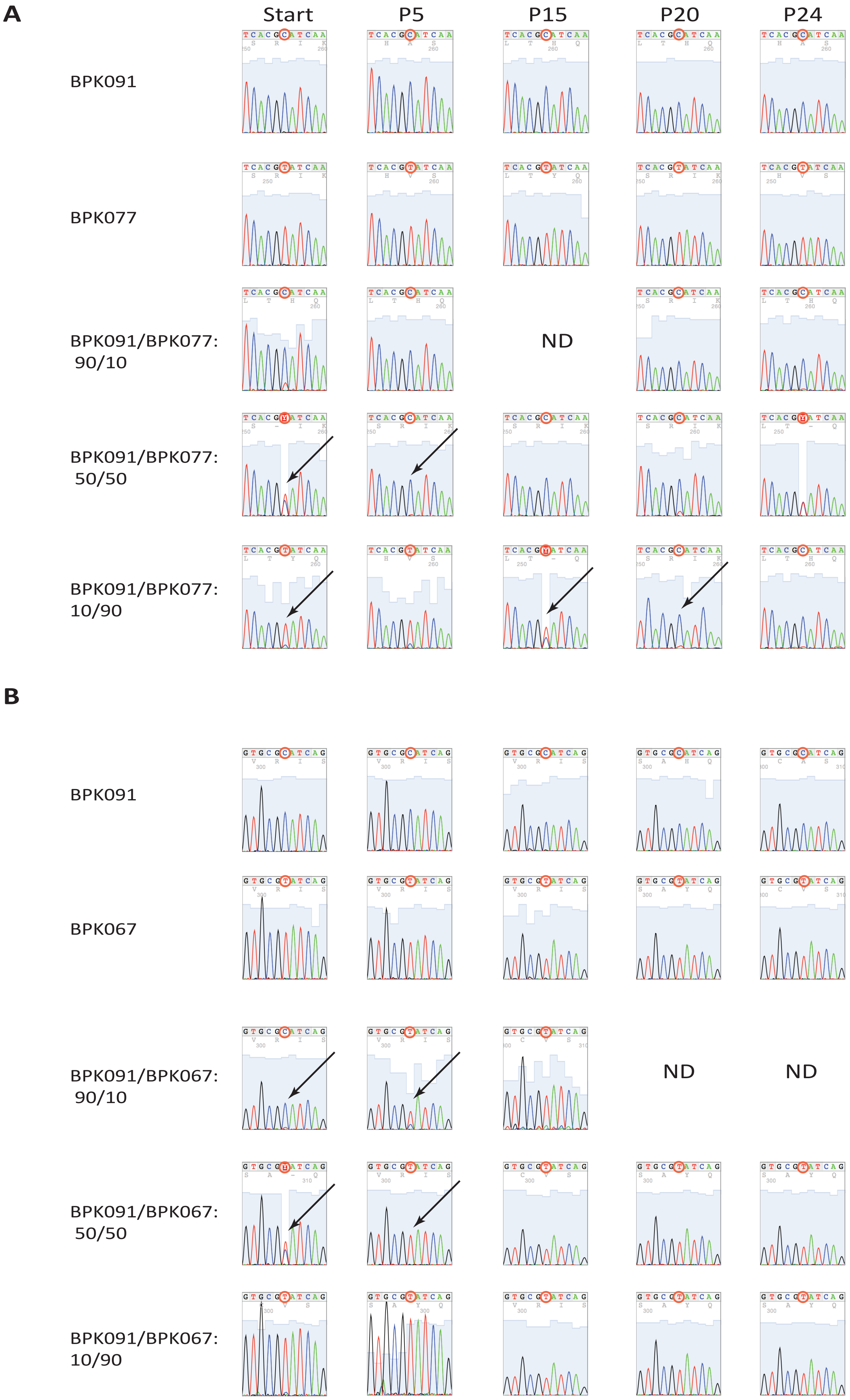


**Fig. S8. Growth competition experiment.** Flasks were inoculated with individual cloned strains or mixtures of two cloned strains in the following ratios: 90/10, 50/50 and 10/90. Cultures were analyzed at the beginning of the experiment (Start), after 5, 15, 20 and 24 passages (P5, P15, P20 and P24, respectively). Diagnostic PCR that allows to distinguish between the two compared strains was used to monitor the presence of each strain in culture. A selected fragment of chromatogram containing the variable nucleotide (labelled with red circle) is shown. Arrows point at the samples were the changes in dominance occurred. **A**. Competition between the BPK091, and BPK077 strains (ISC4, and ISC6, respectively). **B**. Competition between the BPK091, and BPK067 strains (ISC4, and ISC3, respectively).
